# Species and Cell-Type Properties of Classically Defined Human and Rodent Neurons and Glia

**DOI:** 10.1101/212696

**Authors:** Xiao Xu, Elitsa I. Stoyanova, Agata Lemiesz, Jie Xing, Deborah C. Mash, Nathaniel Heintz

**Affiliations:** Laboratory of Molecular Biology, Howard Hughes Medical Institute, The Rockefeller University, 1230 York Avenue, New York, NY 10065, USA; University of Miami Miller School of Medicine, Miami, FL 33136, USA

## Abstract

Determination of the molecular properties of genetically targeted cell types has led to fundamental insights into mouse brain function and dysfunction. Here, we report an efficient strategy for precise exploration of gene expression events in specific cell types in a broad range of species, including postmortem human brain. We demonstrate that classically defined, homologous neuronal and glial cell types differ between rodent and human by the expression of hundreds of orthologous, cell specific genes. Confirmation that these genes are differentially active was obtained using epigenetic mapping and immunofluorescence localization. Studies of sixteen human postmortem brains revealed cell-specific molecular responses to aging, and the induction of a shared, robust response to an unknown external event experienced by three donors. Our data establish a comprehensive approach for analysis of unique molecular events associated with specific circuits and cell types in a wide variety of human conditions.

## Introduction

The ability to genetically target (Gong et al., 2003) and molecularly profile specific cell types (Heiman et al., 2008) has begun to provide insight into essential features of the mammalian brain that were discovered in the founding studies of Ramon y Cajal over a century ago (Ramon y Cajal et al., 1899). It has been established, for example, that each anatomically distinct, classically defined cell type expresses a set of genes that is characteristic (Dougherty et al., 2010; Doyle et al., 2008), that these genes confer properties that are essential for specialized cellular functions (Kim et al., 2008; Nakajima et al., 2014), and that the expression of these genes is dependent on maintenance of cell specific epigenetic states that organize nuclear function (Kriaucionis and Heintz, 2009; Mellén et al., 2012). Application of these powerful new technologies in mouse models has led also to the realization that environmental influences (Heiman et al., 2008; Shrestha et al., 2015), internal physiological cues (Schmidt et al., 2012), and disease causing genetic lesions (Fyffe et al., 2008; Ingram et al., 2016) alter gene expression in affected cell types. Despite the pace of advances in experimental systems, fundamental issues of human brain complexity remain unsolved. It is not known, for instance, how many distinct cell types exist in the human brain, how these cell types vary between individuals or across species, whether the process of brain aging is equivalent between cell types, and why mutations in broadly expressed genes can have devastating consequences in one or a few select cell types.

To address these questions, two main approaches have been taken that enable molecular characterization of cell types without the need for transgenic animals. The first involves gene expression profiling at the level of single cells (Darmanis et al., 2016; Macosko et al., 2015; Thomsen et al., 2016; Zeisel et al., 2015) or single nuclei (Grindberg et al., 2013; Habib et al., 2016; Krishnaswami et al., 2016; Lacar et al., 2016; Lake et al., 2016). These studies have been instrumental in providing an unbiased description of the diversity of cell types and even subpopulations of cell types in the mouse and human brain. However, the profiles generated from these types of studies are known to have high levels of variability, both due to technical challenge of amplifying the signal from a small amount of starting material, and biological causes such as transcriptional bursting (Haque et al., 2017). Additionally, because single cell studies typically take an unbiased approach to profiling the cell types in a tissue, common cell types will be sequenced many more times than rare cell types.

A second approach for molecular characterization has been developed to capture specific cell types, usually by labeling followed by fluorescence activated cell sorting (FACS). Because neurons have complex cellular architecture, and techniques to achieve the single cell suspension required for FACS stress and damage neurons, these studies are typically performed with isolated nuclei instead of whole cells (Matevossian and Akbarian, 2008; Molyneaux et al., 2015). These methods have enabled expression and epigenetic profiling of several cell types during development of both the mouse and from both mouse and the human brain (Ernst et al., 2014; Lister et al., 2013; Molyneaux et al., 2015). However, only a few cell types have been characterized using this approach, primarily due to the difficulty of identifying antibodies to nuclear localized proteins that can label the nuclei from cell types of interest with sufficient specificity to enable sorting.

To overcome this limitation, we have taken advantage of the fact that the endoplasmic reticulum membrane (ER) is contiguous with the nuclear membrane (Hetzer, 2010; Watson, 1955). We reasoned that we could expand the number of antibodies useful for purifying cell-type specific nuclei by targeting ER proteins in addition to nuclear regulatory proteins. Here we demonstrate that antibodies against both ER resident proteins and plasma membrane proteins being translated in the ER can be used for labeling and sorting nuclei from postmortem brains, dramatically expanding the pool of antibodies available for cell specific nuclear isolation. As a proof of principle, we used antibodies against a variety of different proteins to purify nuclei and profile gene expression from the major cell types in the mouse cerebellum. We demonstrate that this approach can be extended readily to cell types from rat and human brain, and that the cell specific nuclear expression profiles are reproducible, cell specific and comprehensive. Comparative studies of data from rodent and human brain reveal a surprising degree of evolutionary divergence of gene expression profiles from classically defined, highly conserved cerebellar cell types (D’Angelo, 2013; Eccles, 1967; Llinas, 1969). The data suggest also that molecular mechanisms of brain aging unfold differently in each human cell type. Finally, they establish that robust molecular phenotypes that might indicate a shared external event can be discovered in control postmortem human brains. We anticipate that correlative studies of human genetic and clinical data with celltype specific expression and epigenetic profiling will enable the usage of postmortem tissue from common brain banks to identify as yet unrecognized molecular characteristics of human brain function and dysfunction.

## Results

### A non-genetic method to specifically isolate nuclei from select cell types

To generate reference nuclear profiles for cell types of interest, we employed transgenic mice expressing an EGFP-L10a ribosomal protein fusion under the control of cell-type specific drivers (Figure S1A) (Doyle et al., 2008; Heiman et al., 2008) for purification of nuclei by FACS (Kriaucionis and Heintz, 2009; Mellén et al., 2012). RNA-seq was used to generate nuclear expression profiles for five different cell types – cerebellar Purkinje cells, granule cells, or Bergmann glia, or corticopontine or corticothalamic pyramidal neurons (Figure S1B, C). Analysis of these data and published data for three cortical cell types (excitatory neurons, PV interneurons, and VIP interneurons, Mo *et al.* (2015)) demonstrated that nuclear RNA expression profiles can both distinguish different cell types and reveal relationships between cell types (Figure S1D). This finding confirms the utility of nuclear RNA profiles for characterization of CNS cell types.

Next we generated expression profiles from non-transgenic brains, several classically defined neuronal and glial cell types that exhibit unique morphology and distribution in the cerebellum (Figure 1A). Candidate antibodies for each cell type were chosen from our previously published translating RNA (TRAP) data (Dougherty et al., 2010; Doyle et al., 2008). To isolate three neuronal cell types (granule, Purkinje, basket) and two glial cell types (astrocytes and oligodendrocytes) from the cerebellum of wild-type mice, we purified and then labeled nuclei (Figure S2A) using antibodies against one transcription factor (OLIG2), one ER resident membrane protein Inositol 1,4,5-Triphosphate Receptor Type 1 (ITPR1), and two plasma membrane proteins Sortilin Related VPS10 Domain Containing Receptor (SORCS3) and Glutamate Aspartate Transporter (EEAT1 or GLAST). To facilitate separation of neuronal from glial nuclei, we used in combination with each of these cell-type specific antibodies an antibody against the splicing protein RBFOX3/NeuN. We also used NeuN to positively select granule cells while removing nuclei from contaminating cell types using a combination of ITPR1, SORCS3, and OLIG2 (Figure 1B). Imaging of nuclei labeled with these antibodies confirmed the expected distribution of the target proteins – while the nuclear factors NeuN and OLIG2 are localized within the nucleus, specifically in euchromatin, the ER resident protein ITPR1 and the plasma membrane proteins SORCS3 and EEAT1 localize to the ER/nuclear membrane and appear as a ring around the nucleus (Figure 1D-E). However, both flow cytometry and imaging of the labeled nuclei yielded a surprising result: although NeuN is not known to label Purkinje nuclei in cerebellar tissue, staining of nuclei using NeuN robustly labels Purkinje nuclei (Figure 1D). This is not due to spectral overlap from the Purkinje specific marker, as ITPR1 localizes to the ER/nuclear membrane while NeuN localizes to euchromatin, both as expected. Although we do observe lower levels of NeuN in labeled basket, astrocyte, and oligodendrocyte nuclei, this result suggests the exposure of NeuN epitopes might be slightly different between isolated nuclei and tissue sections.

**Figure 1.**
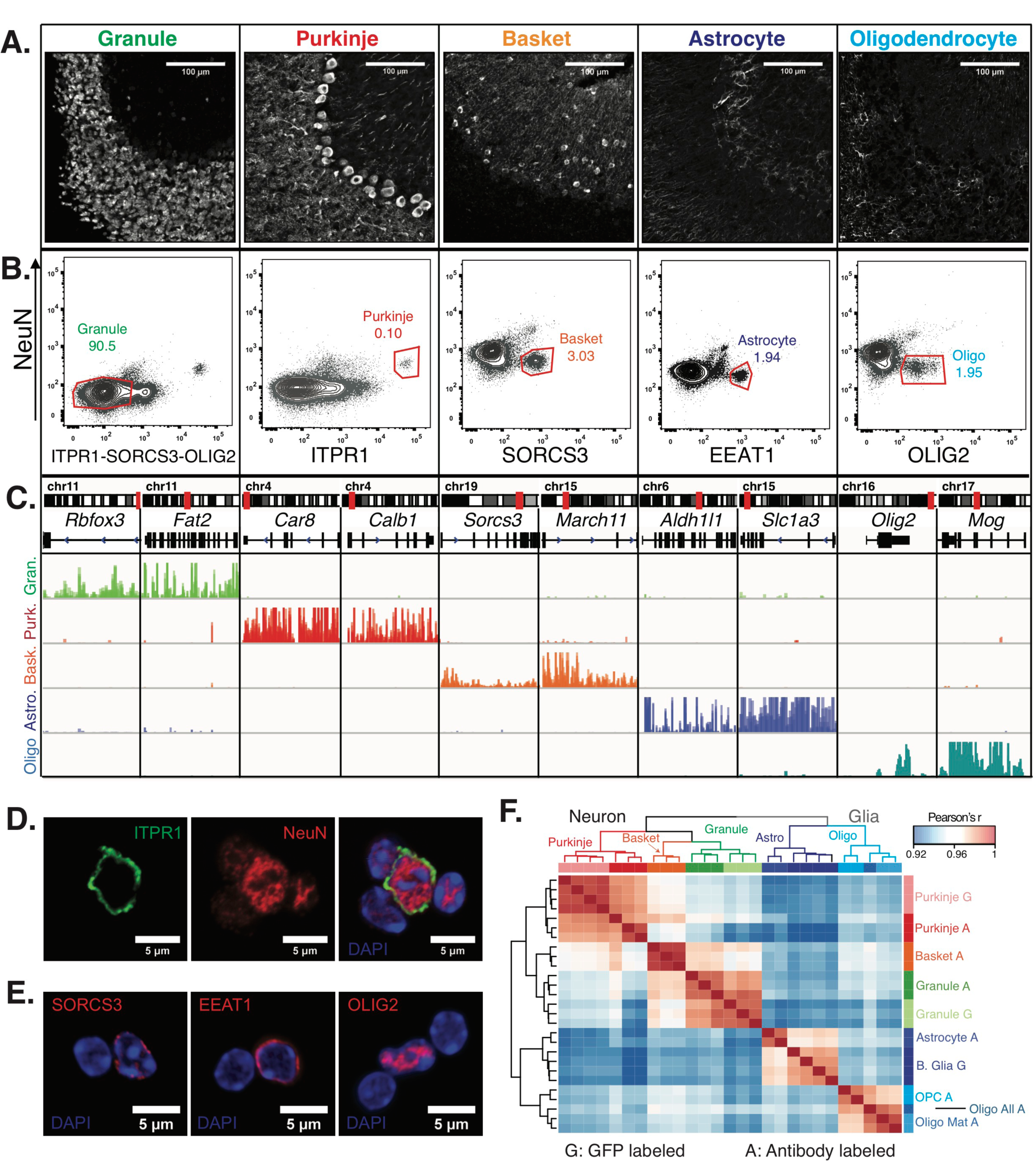
Generation of gene expression profiles for distinct cell types from the cerebella of wild-type mice. **(A)** Immunofluorescence staining of five distinct cell types in the cerebellum. Antibodies used for each cell type: NeuN for granule cells, ITPR1 for Purkinje cells, SORCS3 for basket cells, GFAP labels the cell bodies and process of astrocytes, MOG labels the cell bodies and process of oligodendrocyte. **(B)** Fluorescence activated sorting of stained nuclei from five cell types. Antibodies used for staining are indicated on the x-and y-axes. Percentage of each cell type based on the positive population is indicated. **(C)** Browser view showing examples of gene that are specifically expressed in each of the five cell types. **(D,E)** Examples of labeled cerebellar nuclei using cell-type specific antibodies. Nuclei are counterstained with DAPI, a marker for heterochromatin. **(D)** Nuclei labeled with antibodies against ITPR1 and NeuN. Itpr1, an endoplasmic reticulum membrane protein, is localized at the nuclear membrane, while NeuN, a splicing factor, is localized at euchromatin inside the nucleus. **(E)** Antibodies against the basket cell marker SORCS3 and astrocyte marker EEAT1, two cellular membrane proteins, show labeling of the nuclear membrane. Antibodies against the oligodendrocyte marker and transcription factor OLIG2 show labeling in euchromatin. **(F)** Heatmap showing the pairwise Pearson’s correlation coefficient of GFP- (G) and antibody- (A) sorted nuclei. Hierarchical clustering is performed on the 250 most variable genes across all conditions. See also Figures S1, S2.

### Nuclear RNA profiles generated from sorted nuclei are cell-type specific

We next produced gene expression profiles for each of the five cerebellar cell types by isolating RNA from the sorted nuclei and performing RNA-seq. To evaluate the purity of these profiles, we checked the expression of known markers for each cell type of interest. In all cases, we observed high expression of markers only in the appropriate cell type. For example, the granule cell markers *Rbfox3* (NeuN) and *Fat2* are highly expressed in granule cells and are minimally expressed in the other four cell types (Figure 1C). Next, we evaluated these samples at a genome-wide level by computing the pairwise Pearson’s correlation coefficient (r) between all samples using normalized gene expression (Figure 1F). From this analysis, we could make several observations. First, the correlation between different cell types in the cerebellum is high: r=0.91-0.94. Second, the correlation across biological replicates is very high: r=0.98-1. Third, when we compared to the reference gene expression profiles from the EGFP-L10a labeled nuclei from granule cells, Purkinje cells, and Bergmann glia, we observed very high correlations between the genetically defined cell types and antibody defined cell types: r=0.98 for granule cells and r=0.99 for Purkinje cells; r=0.96-0.97 between Bergmann glia purified from *Sept4* EGFP-L10a animals and astrocytes purified via antibody labeling (Figure 1F). Since Bergmann glia are a sub-population of cerebellar astrocytes, it makes sense for these datasets to have slightly lower correlations. Finally, when we perform hierarchical clustering, the samples cluster according to known biology. Thus, neuronal and glial samples separate from each other in the first split. The five cell types then separate from each other regardless of whether the nuclei were purified using genetic labeling (G) or antibody labeling (A). Taken together, our results demonstrate that that fluorescence activated nuclear sorting allows cell-type specific gene expression profiling without the requirement for transgenic animals.

### Antibody-based sorting enables the isolation and identification of subpopulations of cell types

In the FACS isolation of oligodendrocytes, we noticed that the OLIG2+ population could be subdivided based on its levels (Figure S2B). By flow cytometry, we found that approximately 20% of all OLIG2+ nuclei have high levels OLIG2 (High) while 80% have lower levels (Low). To determine whether these two populations form distinct subpopulations of oligodendrocytes, we isolated and analyzed them individually using RNA-seq. We found that both populations express general oligodendrocyte markers such as OLIG2, but that only OLIG2+ High nuclei express genes such as *Pdgfra* and *Cspg4* that mark oligodendrocyte precursor cells (OPCs) (Figure S2C). We conclude that the OLIG2+ Low population contains mature oligodendrocytes, while OLIG2+ High nuclei contains a mix of mature oligodendrocytes and immature OPCs. This is consistent with previous studies demonstrating that OLIG2 levels are higher in oligodendrocyte precursors compared to mature oligodendrocytes (Hayashi et al., 2011; Kitada and Rowitch, 2006; Kuhlmann et al., 2008). Furthermore, hierarchical clustering of nuclear RNA-seq profiles from all cerebellar cell types indicates that the oligodendrocyte lineage splits into two groups, mature oligodendrocytes and OPCs, with the set of all oligodendrocytes clustering with mature oligodendrocytes (Figure 1F). These results indicate that fluorescence activated sorting can also be used to identify subpopulations of cell types. In succeeding text and figures, these two subpopulations are analyzed separately and mature oligodendrocytes referred to simply as oligodendrocytes.

### Comparison of expression data from cell type specific populations of nuclei to single nuclei

The application of population based, cell type specific technologies for expression profiling in the human brain provides an important complement to ongoing surveys of CNS cell types using single cell analysis. In particular, the ability to isolate RNA from a large number of defined nuclei ensures deeper and more comprehensive data than is currently available from single cell studies. To demonstrate that this is the case, we compared our cerebellar glial datasets with hippocampal glial single nucleus sequencing results from Habib et al. (2016). Cumulative distribution function plots (Figure S2D) indicate that while the distribution of normalized reads per gene is similar between EGFP+ sorted Bergmann glia data from transgenic animals and antibody sorted astrocyte, oligodendrocyte, and OPC data from wild-type animals, the single nuclei data have a very high number of genes with zero reads. This is not simply the result of sequencing depth (Figure S2E) because even when we subsample our data to equivalent depths (e.g. ~1M reads) the population based data reveals expression of 10-12,000 genes whereas single cell data allow detection of 5-6,000 expressed genes. Additionally, expression of general or cell-type specific genes is more variable across single nuclei samples than in our biological replicates for the same cell types (Figure S2F). While the variability present in single nuclei data may be desirable for addressing some biological questions, we believe that the reproducibility evidenced by highly correlated biological replicates can present important advantages for detection of cell specific molecular phenotypes associated with human pathophysiology.

### Antibody-based sorting of cell types is easily transferrable to other species

An important reason for development of this approach was to enable cell-type specific profiling in species where transgenic strains are not readily available. As a first test of this property, we chose the rat because it is a well-established model organism for behavioral neuroscience. Nuclei were isolated from rat cerebellum, fixed and stained using the same antibodies that we used in mouse, and the results were analyzed on a flow cytometer (Figure 2A). The staining profiles of rat nuclei resembled those of mouse (Figure 1A), although there were differences. For example, in the rat cerebellum, NeuN staining resulted in more distinct positive and negative nuclear populations than was evident in the mouse cerebellum. As a result, additional populations of nuclei were revealed, and these could be isolated and characterized by RNA-seq. It was possible, for example, to isolate nuclei from basket cells, astrocytes, and oligodendrocytes by staining with antibodies against SORCS3 and NeuN. This flexibility allowed us to purify all six cell types using only four types of staining (Figure 2A). For cell types that were isolated from more than one type of staining, we chose the population that by RNA-seq was most enriched for known markers (*) and depleted for markers of other cerebellar cell types.

**Figure 2.**
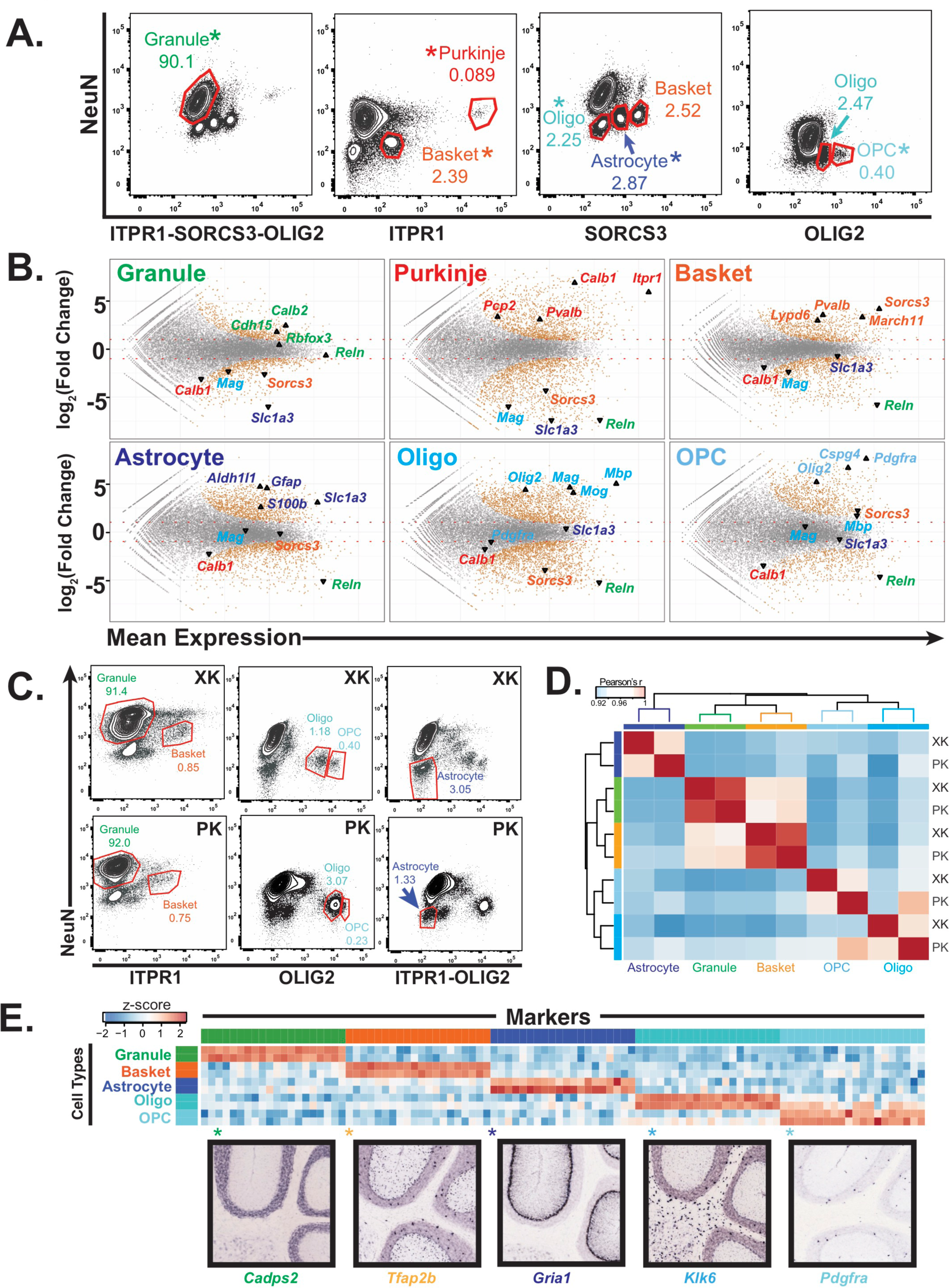
Generation of gene expression profiles for distinct cell types from rat and human cerebella. **(A)** Fluorescence activated sorting of stained nuclei from six cell types in the rat cerebellum. Antibodies are indicated on the x and y axis. When a cell type can be isolated from more than one staining scheme, the population used for downstream analysis is indicated with (*). Percentage of population in each gate is indicated. **(B)** Differential expression analysis of antibody sorted nuclei for six cell types compared to unsorted nuclei from the rat cerebellum. Known markers for each cell type are highlighted: granule (*Cdh15*, *Calb2*, *Rbfox3*, *Reln*), Purkinje (*Pcp2*, *Pvalb*, *Cabl1*, *Itpr1*), Basket (*Lypd6*, *Pvalb*, *Kit*, *Sorcs3*), Astrocyte (*Aldh1l1*, *Gfap*, *S110b*, *Slc1a3*), all oligodendrocytes (*Olig2*), mature oligodendrocyte, (Labeled Oligo: *Mag*, *Mog*, *Mbp*), oligodendrocyte precursor cells, (Labeled OPC: *Cspg4*, *Pdgfra*). **(C)** Fluorescence activated sorting of granule, basket, astrocyte, mature oligodendrocyte, OPC nuclei from the cerebellum of two human samples (XK and PK). Percentage of population in each gate is indicated. **(D)** Heatmap showing Pearson’s correlation coefficient between human samples. Hierarchical clustering is performed using the 250 most variable genes across samples. **(E)** Heatmap showing the 20 most specific genes for each human cell type as identified by the Specificity Index algorithm. Rows: sorted nuclei from XK, PK samples; columns: genes enriched in each cell type. Color represents the z-score of gene expression compared to all samples. Lower panels show examples of gene expression from the Allen Mouse Brain Atlas for genes identified as highly specific based on the Specificity Index analysis of human cell types. See also Figure S3.

To determine the group of genes that are specific to each of the rat cell types, we performed differential gene expression analysis comparing the nuclear expression profile for each cell type to unsorted cerebellar nuclei (Figure 2B). As expected, we observed a general enrichment for known markers for each cell type and a depletion for markers for all other cell types. In addition, our data identified 700 to 1,600 expressed genes that are significantly enriched in each of the rat cell types. Genes that are most specific for each cell type were identified using an updated version of the specificity index algorithm (Dougherty et al., 2010) (Figure S3A, Table S3). An advantage of this approach is that it can identify specifically expressed genes even in cell types that are very abundant in the region of interest (e.g. cerebellar granule cells). To validate both our cell-type specific rat data and the updated specificity index algorithm, we examined the expression of genes identified as being specific to each cell type using the Allen Mouse Brain Atlas *in situ* hybridization database (Figure S3A). For the top 20 most specific genes for all cell types, we observe labeling specifically in the defined cell type on average 93% of the time.

### Antibody-based sorting of cell types can be applied to post-mortem human tissue

To determine whether antibody based nuclear sorting can be used productively for analysis of cell types in the human brain, we acquired post-mortem human cerebellar tissue from two donors (codes XK and PK), then isolated and stained nuclei using the cell-type specific antibodies defined previously. Although the profile of stained human cerebellar nuclei is generally similar to rodent, there are differences (Figure 2C). For example, the conditions used to sort granule and basket neurons, astrocytes, oligodendrocytes, and OPCs were directly transferrable to human postmortem tissue, whereas those used to sort mouse and rat Purkinje cell nuclei were not successful for isolation of human Purkinje cell nuclei, possibly because while Purkinje cells are already rare in mouse and rat (0.1% in mouse, 0.05% in rat; Figures 1B, 2A), they have been reported to be even more rare in human (Korbo and Andersen, 1995). We also observed slight differences in staining between the two postmortem human samples: in XK the oligodendrocyte and OPC populations are well separated, whereas they are more difficult to distinguish in the sample PK. To assess whether these differences might prevent cell type specific analysis of human postmortem samples, we profiled gene expression in nuclei isolated from five cell types (granule, basket, astrocyte, oligodendrocyte, OPC) in these two individual brains using RNA-seq. Examination of known markers from each of these cell types established that relevant cell specific markers are enriched in the data for each cell type, and depleted from the other cell types (Figure S3B). Calculation of pairwise Pearson correlation coefficients across the ten datasets revealed that the two neuronal cell types (granule and basket) were highly correlated between samples (r=0.99 for both), while the glial data had slightly lower correlations (r=0.96-0.97). Hierarchical clustering resulted in separation of the samples by cell type, indicating that the variation across cell types is stronger than that between individual samples (Figure 2D). Finally, we applied the specificity index algorithm to these data to identify specifically expressed genes in each human cell type (Figure 2E). When we examined the genes that are most specific, we noted that many of these genes were also expressed highly in the same cell type in mouse and rat, but noted the presence of some genes that appeared to be human-specific in each cell type (Table S3). We next decided to explore this difference in greater detail.

### Cross-species comparison reveals cell-type and species specific genes

Given the high quality of the cell specific expression profiles described above, we were next interested in addressing directly the extent of gene expression conservation and divergence between species in these well-defined, classical cerebellar cell types. To ensure that our results reflected altered expression rather than differences in annotation between species, we limited our analysis to highconfidence 1:1 orthologous genes between mouse, rat and human (Figure S4A, methods). To explore the similarities and differences across species, we used hierarchical clustering to determine the relationships between samples. Interestingly, we found that when we clustered using all genes, the samples resolved primarily by species (Figure S4B). However, when we performed clustering using only the 250 most variable genes across samples, they cluster primarily by cell type (Figure 3A). To further investigate this result, we computed the pairwise Pearson correlation coefficients across all samples (Figure S4C). We found that the range of correlation coefficients between the samples for the same cell type across species is similar to that for cell types isolated from the same species, suggesting that both cell type and species are important properties of the expression profiles. Principle components analysis provided additional support for this finding (Figures S4D-F). Examination of the first eight principle components (PC1−8), which collectively account for 90.5% of the variability across all samples, revealed that PC1 and 3 categorize samples by cell type, while PC2, 6, and 8 split samples by species. The other principle components organize samples by both cell type and species. Because PC1 and 3 account for 47% of the variability across samples while PC2, 6, and 8 account for only 22% of variability (Figure S4D), we conclude that the most important differences between the profiles result from cell-type specific expressed genes that are shared among species. However, it is apparent that genome-wide quantitative differences in expression profiles between species must also be considered when assessing the fine-tuned functional properties of a given cell type in different species. We note that while we have tried to minimize the effect of annotation differences between species by limiting our analyses to high-confidence 1:1 orthologous genes, it is possible that some of the species differences that we observe can be attributed to differences in annotation. This may explain why hierarchical clustering using all genes results in clustering by species while clustering with the most variable genes results in clustering by cell type. Finally, to gain insight into cell-type specific functions that have changed across species, we calculated the specificity index for all genes in a given species. We next compared the rank of the 100 most specific genes for each cell type in mouse with their ranks in rat or human (Figure S4G, H, Table S3). We found that in general, genes that are highly specific in mouse are also highly specific in rat and human, but with many exceptions. As expected, the specificity rank was more conserved between mouse and rat than mouse and human.

**Figure 3.**
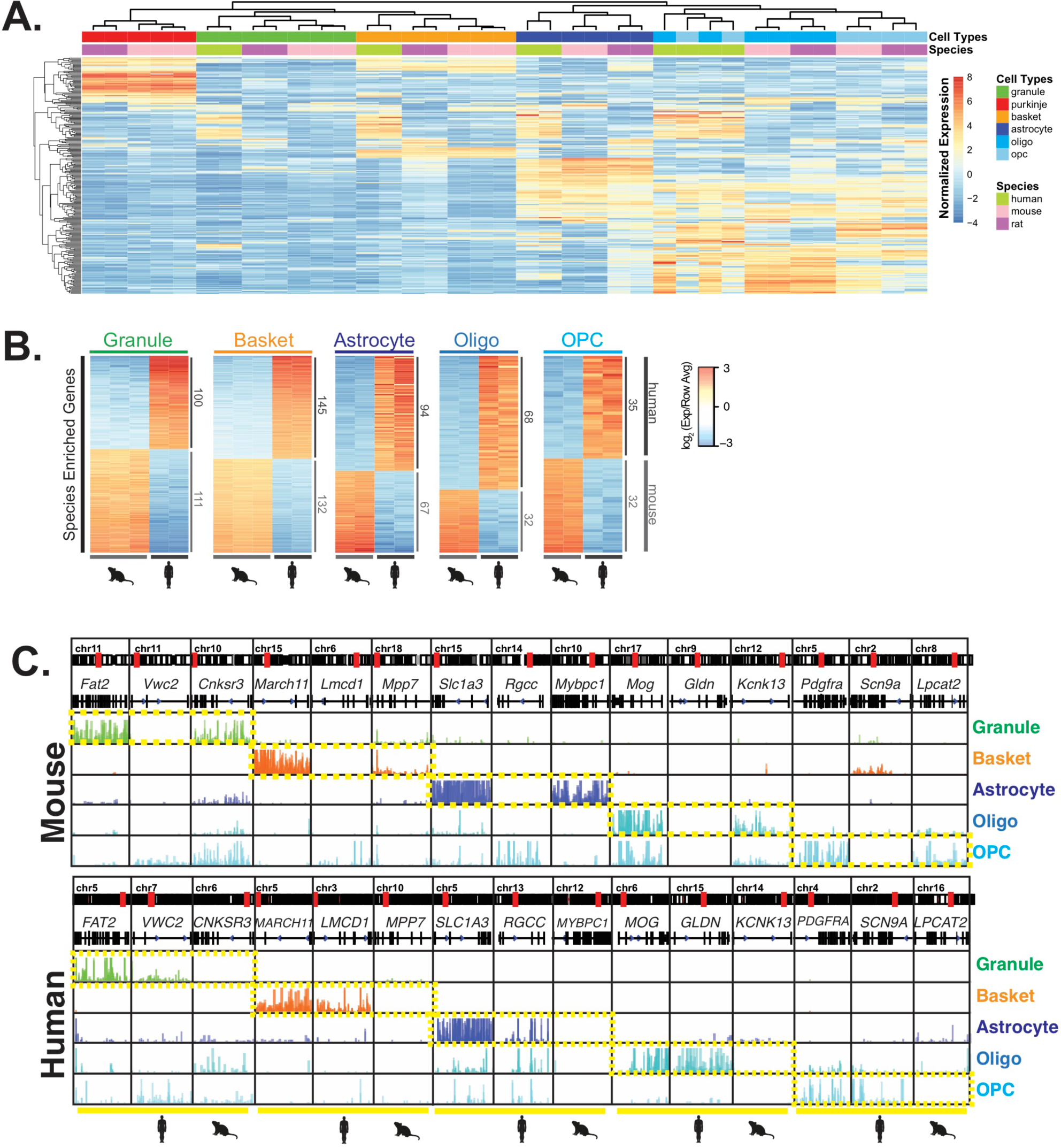
Comparative analysis of gene expression across species reveals cell-type and species specific differences. **(A)** Heatmap showing normalized expression of the 250 most variable genes across all samples. Hierarchical clustering is performed using these genes and reveals that samples cluster primarily by cell type and secondarily by species. **(B)** Heatmap showing mouse- and human-enriched genes for each cell type, excluding any genes that are mouse- or human-enriched across all cell types. The number of genes that are significantly human or mouse enriched is indicated. **(C)** Browser view showing for each cell type, an example of a shared marker gene, a human-enriched gene, and a mouse-enriched gene. See also Figures S4, S5.

To explore further the extent to which cell-type specific genes change across species, we focused our comparative analysis on the data from mouse and human because the annotation of the rat genome is less complete. By excluding rat genes, the number of high confidence orthologs that could be used for comparison increased from 11,443 to 14,273 (Figure S5A). To focus on differentially expressed genes that are most likely to impact cellular function, we selected for genes that passed a stringent adjusted p-value cutoff of < 10e-5, that changed by at least 4-fold between species, and that are expressed at significant levels in each cell type (average number of normalized counts/DEseq2 baseMean > 400 and log_2_(fpkm) > 0). Using these criteria, we found between 119 and 410 genes that are mouse- or human-enriched in these cell types (Figure S5B). To identify cell-type and species differences rather than general differences between mouse and human, we removed from the analysis genes that are differentially expressed between mouse and human in all cell types analyzed (adjusted p < 10e-5, fold change > 2). Filtering in this way yielded between 67 and 277 cell-type and species specific differentially expressed genes that were approximately equally distributed between those enriched in mouse and those enriched in human (Figure 3B, S5B, Table S4). Examples of shared and species specific expressed genes are shown in Figure 3C. Interestingly, expression of these genes in rat most often resembles expression in mouse, although there are exceptions (Figure S3C). Gene Ontology (GO) analysis was performed for all ten categories of species-specific genes to understand whether their expression might reflect changes that occur independently, or be due to transcriptional programs that are altered between species. Six of these gene sets do not have significantly associated GO categories suggesting species differences in these cell types contribute to a wide range of pathways and functions. GO analysis of human-enriched genes from astrocytes revealed weak enrichment for genes involved endocytosis regulation. GO analysis of mouse-enriched genes from astrocytes revealed weak enrichment for genes with phosphoric diester hydrolase / phospholipase activity, while mouse-enriched genes from basket and granule cells revealed enrichment for genes with transporter activity (Table S6).

To confirm the presence or absence of some of these species and cell-type specific genes (Figure S6A) in the mouse brain, we examined publicly available gene expression data from the Allen Mouse Brain Atlas and GENSAT Project (Gong et al., 2003; Lein et al., 2007) (Figure S6B, C). These data demonstrate expression in the granule layer of the cerebellum for the mouse-enriched genes *Cnksr3*, *Ece1*, and *Pde1c* (Figure S6B), and the absence of expression of the human-enriched genes *Ccdc175*, *Clvs2*, *Vwc2*, and *Pde1a* (Figure S6C). Interestingly, while *Clvs2* and *Pde1a* are absent from the cerebellum, they are expressed in other regions in the mouse brain, suggesting that alterations in cell specific regulatory sequences may underlie these species differences (Carroll, 2005; 2008).

To provide further confirmation of these data, we took advantage of the abundance of granule cell neurons (>90%) relative to all other cell types in the cerebellum (Figure S1B, 1B, 2A, 2C, Table S1). This allowed two types of validation that could not be obtained for less abundant cell types: first, in addition to assaying the expression of nascent nuclear RNAs, we could examine the expression of mature cytoplasmic RNAs; second, given the high fraction of granule cell genomes present in cerebellar nuclear preparations, chromatin accessibility could be assessed using unsorted cerebellar nuclei. Since previous studies have demonstrated that ATAC peaks are typically found in the promoters and gene bodies of expressed genes and severely reduced or absent from genes that are not expressed (Buenrostro et al., 2013; Mo et al., 2015; Su et al., 2017), we collected ATAC-seq data from mouse, rat and human cerebellar nuclei to confirm the expression data. As expected, protein coding genes that are identified as expressed in our data are accompanied by the presence of spliced cytoplasmic mRNAs, and ATAC peaks are evident in their promoters and gene bodies. For example, in the locus containing *GPR135* [1], *L3HYPDH* [2], *JKAMP* [3], *CCDC175* [4], and *RTN1* [5], all five genes are expressed in human granule cells as revealed by nuclear and cytoplasmic gene expression and the presence of ATAC peaks (Figure 4A, bottom). However, in mouse by nuclear and cytoplasmic gene expression, *Rtn1* [5] is expressed, *Ccdc175* [4] is not expressed, and *Jkamp* [3], *L3hypdh* [2], and *Gpr135* [1] are expressed at much lower levels relative to their human orthologs. At this locus in mouse, ATAC peaks remain present in the promoters or gene bodies of the expressed genes *Rtn1*, *Jkamp*, *L3hypdh*, and *Gpr135*, but are completely missing over the promoter and gene body of nonexpressed gene *Ccdc175* (Figure 4A, top).

**Figure 4.**
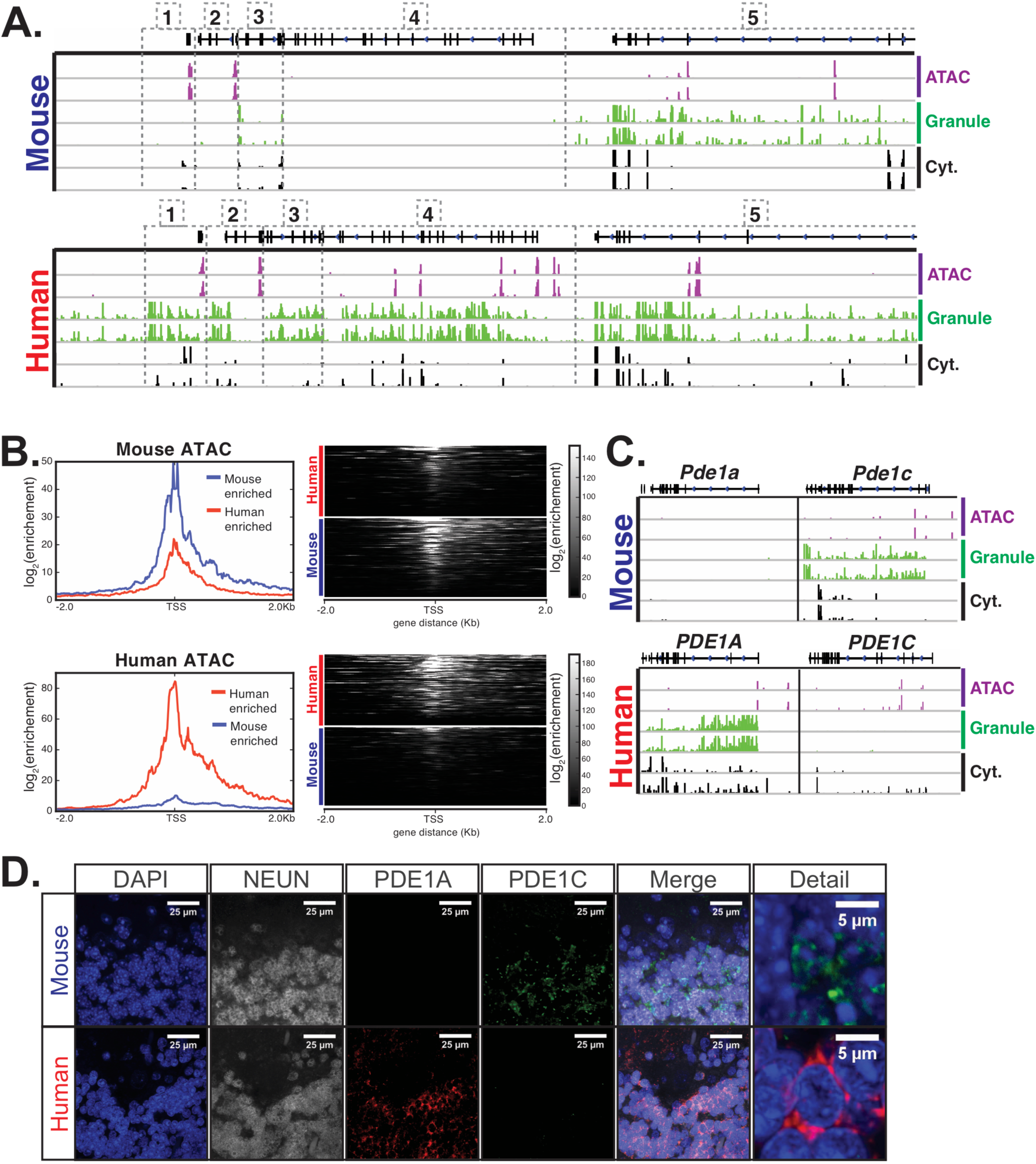
Epigenetic and immunofluorescence validation of gene expression differences between mouse and human cerebellar granule cells. **(A)** Browser views showing a homologous region of approximately 300kb from chr12 in mouse and chr14 in human. Genes located in this region are: [1] *Gpr135* [2] *L3hypdh* [3] *Jkamp* [4] *Ccdc175* [5] *Rtn1*. The three types of tracks shown are: ATAC-seq DNA accessibility from total cerebellum (purple), nuclear RNA levels from granule cells (green), and cytoplasmic RNA levels from total cerebellum (black). Two biological replicates are shown for all tracks. All five genes in the locus are strongly expressed in human granule, but only *Rtn1* is similarly expressed in mouse granule cells. *Ccdc175* is not expressed in mouse, as evidenced by the lack of granule cell and cytoplasmic RNA and the absence of DNA accessibility sites in its promoter and across its gene body. The other genes – *Jkamp*, *L3hypdh*, and *Gpr135* – are expressed in mouse, as supported by the presence of promoter DNA accessibility sites, but at lower levels compared to human. **(B)** Analysis of ATAC-seq DNA accessibility assay from mouse (top) or human (bottom) from unsorted cerebellar nuclei. Granule cell nuclei are >90% of the total in both mouse and human samples. Left: metagene analysis showing the median log_2_ fold enrichment of reads from the promoter regions of 100 human-enriched or 111 mouse-enriched granule cell species specific genes. Right: density plot showing enrichment of ATAC peaks over the promoter of all 211 genes. **(C)** Example of gene usage switching between species. Expression of two *Pde1* family members – *Pde1a* and *Pde1c* – in granule cells from mouse and human. Mouse granule cells express *Pde1c* but do not express *Pde1a,* as evidenced by the presence of DNA accessible sites, granule cell nuclear RNA, and total cerebellar cytoplasmic RNA for *Pde1c* but not *Pde1a*. In contrast, granule cells from human cerebellum express high levels of *PDE1A,* as demonstrated by the presence nuclear transcripts, cytoplasmic RNA, and ATAC peaks (purple) indicating DNA accessibility in the promoter and gene body. PDE1C is expressed in human cerebellum at much lower levels, as indicated by all three measures of active gene expression. **(D)** Immunofluoresence confirming the expression of PDE1A and PDE1C in mouse and human cerebellar slices. In mouse, PDE1C is present specifically in granule cells. PDE1A is not expressed above the background of the assay. In human cerebellum, PDE1A is evident specifically in granule cells, and background labeling is observed for PDE1C. NeuN is a marker for granule cells. Staining was performed at least two times each using sections from two separate mice and two human donors. Images shown are representative of all data collected See also Figures S5, S6.

We next analyzed ATAC reads over the promoters of the 100 human-enriched and 111 mouse-enriched granule cell specific genes that we identified by differential gene expression analysis. We found that in mouse, most of the 111 mouse-enriched genes have strong ATAC signal over the promoter region, while only a few of the 100 human-enriched genes have strong promoter ATAC signals (Figure 4B – top). In human, the reverse is true – most of 100 human-enriched genes have strong promoter ATAC signals compared to only a few of the 111 mouse-enriched genes (Figure 4B – bottom). These data both confirm the species differences detected by gene expression analysis and demonstrate that species specific expression for both mouse and human granule cells is accompanied by changes in chromatin accessibility, as revealed by ATAC analysis of the region surrounding the transcription start site (Figure 4B).

While examining the genes that differ in granule cells between mouse and human, we noted an interesting example that involves changing expression of alternative family members. The *Pde1* gene family contains the three calcium- and calmodulin-dependent phosphodiesterases *Pde1a*, *Pde1b*, and *Pde1c*. These enzymes catalyze the hydrolysis of cAMP or cGMP, and they play important roles in modulation of neuronal activity. Our data demonstrate that mouse granule cells mouse express only *Pde1c*, whereas granule cells from human express high levels of *PDE1A* and low levels of *PDE1C* (Figure 4C). Our results match previous findings that *PDE1A* is expressed in the human but not mouse cerebellum (Loughney and Ferguson, 1996; Loughney et al., 1996; Sonnenburg et al., 1993). To further confirm this result, we assayed expression PDE1A and PDE1C in mouse and human cerebellar sections by immunofluorescence (Figure 4D). We found, consistent with the RNA-seq and ATAC data, that both PDE1 members are primarily localized in the granule layer, with PDE1C expression predominant in mouse and PDE1A expression predominant in human cerebellum (Figure 4D). Since PDE1A preferentially hydrolyzes cGMP while PDE1C hydrolyzes both cAMP and cGMP with equal efficiencies (Takimoto, 2009), these differences may result in slightly different biochemical consequences in mouse and human cerebellum.

Taken together, these data demonstrate that nuclear RNA-seq data can be employed to accurately assess cell-type specific gene expression in rodent and human brain, and they allow several important conclusions regarding gene expression in the nervous system. First, gene expression in the brain is highly conserved in specific cell types across species. Accordingly, most abundant and cell specific markers of well characterized cell types are shared between rodent and human brain. Second, despite this shared identity, there are a significant number of genes in each cell type whose expression is not conserved between species. This group of genes does not generally conform to known GO categories, and in many cases, they are expressed in other brain regions. Third, cell type and species specific expression can reflect altered regulation of adjacent genes within a locus, or selective expression of functionally related yet distinct members of a given gene family. Although the consequences of these gene expression changes will have to be interrogated in future studies, our data demonstrate that important differences in gene expression in specific CNS cell types occur between species, and suggest that they may result in functionally important differences in the biochemical functions of even the most classical rodent and human cell types.

### Nuclear profiling of three cell types isolated from 16 post-mortem human brains

Given the genetic heterogeneity of human populations, the variations in human tissue processing times, and the difficulties cited in previous studies of total RNAs isolated from human tissue (McCall et al., 2016; Webster, 2006), we were interested in determining sources of variability across samples from different individuals. To better understand the source of this variability, we analyzed cell-type specific gene expression in an additional 14 human brain samples. In total, we obtained samples from 16 control donors, roughly equally split between genders (7 males and 9 females), and across ages (4 individuals each in the following age groups: 20-30, 40-50, 60-70, 80-90 years) (Figures 2C, 5A, Table S2). We prepared nuclei from all 16 samples and stained them using antibodies against ITPR1 and NeuN as this staining strategy allowed us to purify nuclei from three different cell types in a single sort: granule cells, basket cells, and total glia (Figure 2C, 5A). We noticed that, although the staining pattern varied slightly between samples, we could easily distinguish the ITPR1-NeuN+ population containing granule cells, the ITPR11+ NeuN-population containing basket cells, and the ITPR1-NeuN-population containing total glia. The only exceptions were the glial populations from samples WC and WO in which the ITPR1-NeuN-population did not separate well from the ITPR1-NeuN+ population (Figure 5A). Gene expression analysis of these two samples revealed that these sorts failed to effectively separate granule cells from glia, and they were excluded from further analysis. Consequently, our analysis included 46 human datasets: 16 from granule cells, 16 from basket cells, and 14 from glia. As shown in Figure 5B, RNA-seq analysis of these datasets demonstrate that they are highly pure, as each sample is enriched for the appropriate markers for that cell type, and depleted for markers of other cerebellar cell types. As expected, only samples from females express the X-inactive specific transcript (*XIST*), while only samples from males express Y chromosome genes such as *KDM5D*. Additionally, pairwise Pearson correlation coefficients between biological replicates for each cell type are between 0.98 and 1 (Figure 5C). These data demonstrate that, despite small differences between individuals in the abundance of each cell type or the recovery of nuclei during preparation and sorting, the cell-type specific human expression data obtained by nuclear sorting is highly reproducible.

**Figure 5.**
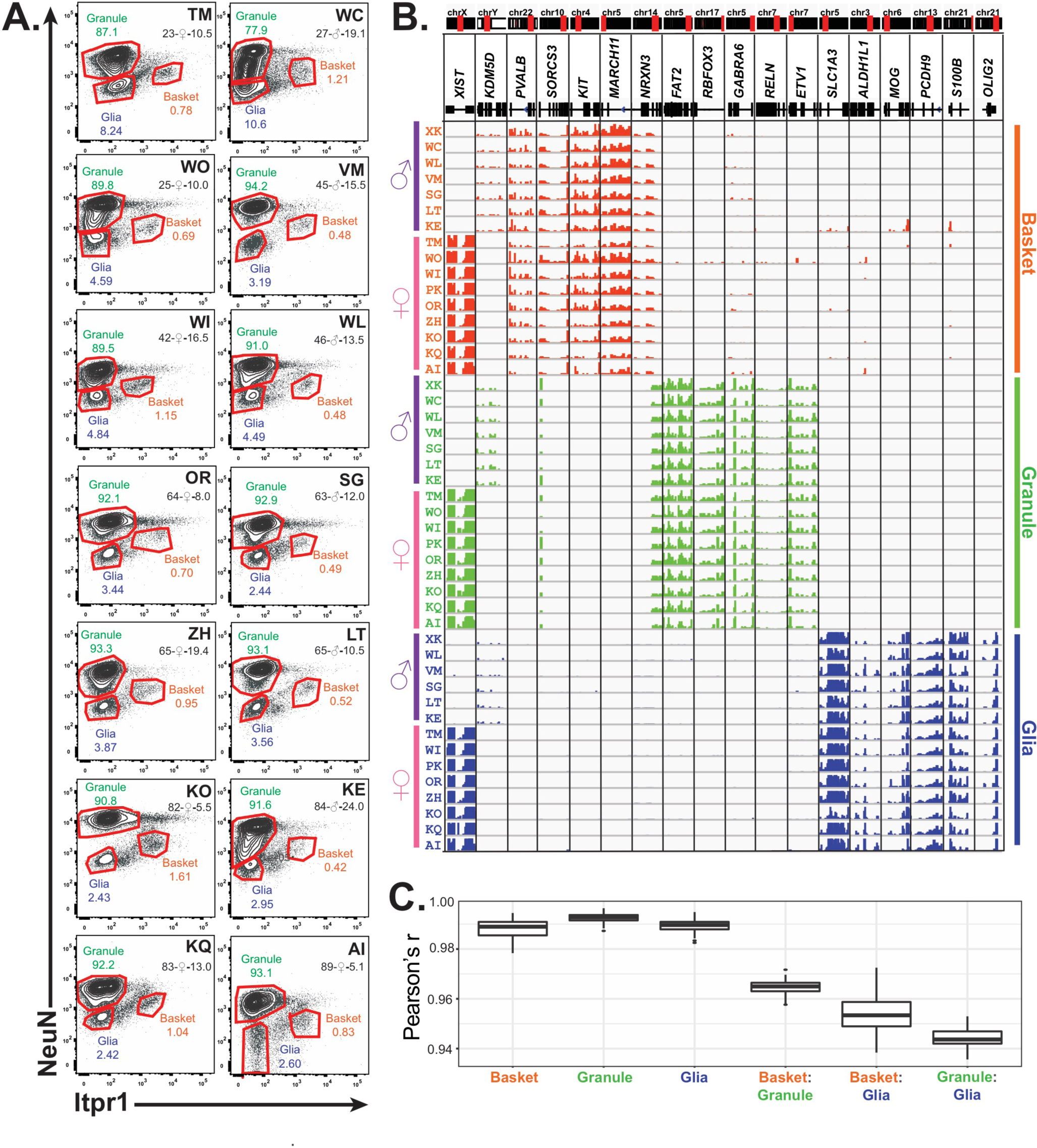
Profiling of three cell types in 16 human cerebellar samples. **(A)** Fluorescence activated nuclear sorting of three cell types from the cerebellum of 14 individuals. Percentages of each cell type in different individuals is shown. Each sample is identified by a two-letter code. Also shown are the age, gender, and autolysis time for each sample. **(B)** Browser view showing gene expression for gender and cell-type markers across 16 individuals (14 from A and two from Figure 4). For glia, samples from WC and WO were excluded due to granule cell contamination. **(C)** Boxplot showing pairwise Pearson’s correlation coefficient within cell types and between cell types for different individuals.

### Factors that impact human cell-type specific gene expression

We next wanted to investigate the relationship between human cell-type specific gene expression and biological and clinical factors that might influence cellular function. Although the cerebellar samples characterized here were obtained from normal controls, differences in gender, age, and the time between death and tissue preservation (autolysis time or postmortem delay interval) might influence the expression data we obtained. We first investigated the effect of autolysis time as a potential source of technical variability that can complicate biological interpretations of the data. We found that only a few genes significantly change with autolysis time: 12 across all samples, four in granule cells, one in basket cells, and one in glia (Figure S7A, Table S5). Examination of these genes revealed that only one, Kif19 in glia, changed linearly with autolysis time (Figure S7C). The remaining genes appear to increase in only a few samples with intermediate autolysis times (Figure S7B). These data confirm an earlier report that autolysis has a relatively minor effect on expression data obtained from post-mortem human tissue (Gupta et al., 2012).

### Gender

To examine the effect of gender on our data, we compared gene expression in samples from males and females and identified genes that are differentially expressed (Figure S7A, Table S5). As expected, we found that the majority of gender-specific genes resided on the X or Y chromosomes, with expression of X-chromosome genes enriched in female samples and Y-chromosome genes enriched in male samples. Of the 27 genes exhibiting gender-specific differential expression (adjusted p-value < 0.01, baseMean > 50), 7 are located on the X chromosome, 14 are on Y, and 6 on autosomes. Interestingly, gender-specific genes can be more differentially expressed in some cell types than in others. For example, we identified Glycogenin 2 Pseudogene 1 (*GYG2P1*) as a male-enriched gene across all samples, although it is clearly more male-enriched in basket cells than granule cells and glia (Figure 6A). This is not due to mismapped reads as a result of duplication, as Glycogenin 2 (*GYG2*) is expressed only in glia and does not exhibit gender-specific expression (Figure S7D). Examination of the Y chromosome locus containing GYG2P1 reveals male-specific basket cell expression across a large region, including GYG2P1 and approximately 300kb downstream (Figure 6B). In contrast, the nearby genes TTY15 and USP9Y are also male-enriched but they are not cell type specific. Genes that exhibit gender and cell-type specific expression in both glia (Figure S7E) and granule cells are also evident (Figure S7F). While the functional significance of these changes is unknown, our results suggests another potential source of sexually dimorphic, cell specific function in the mammalian brain.

**Figure 6.**
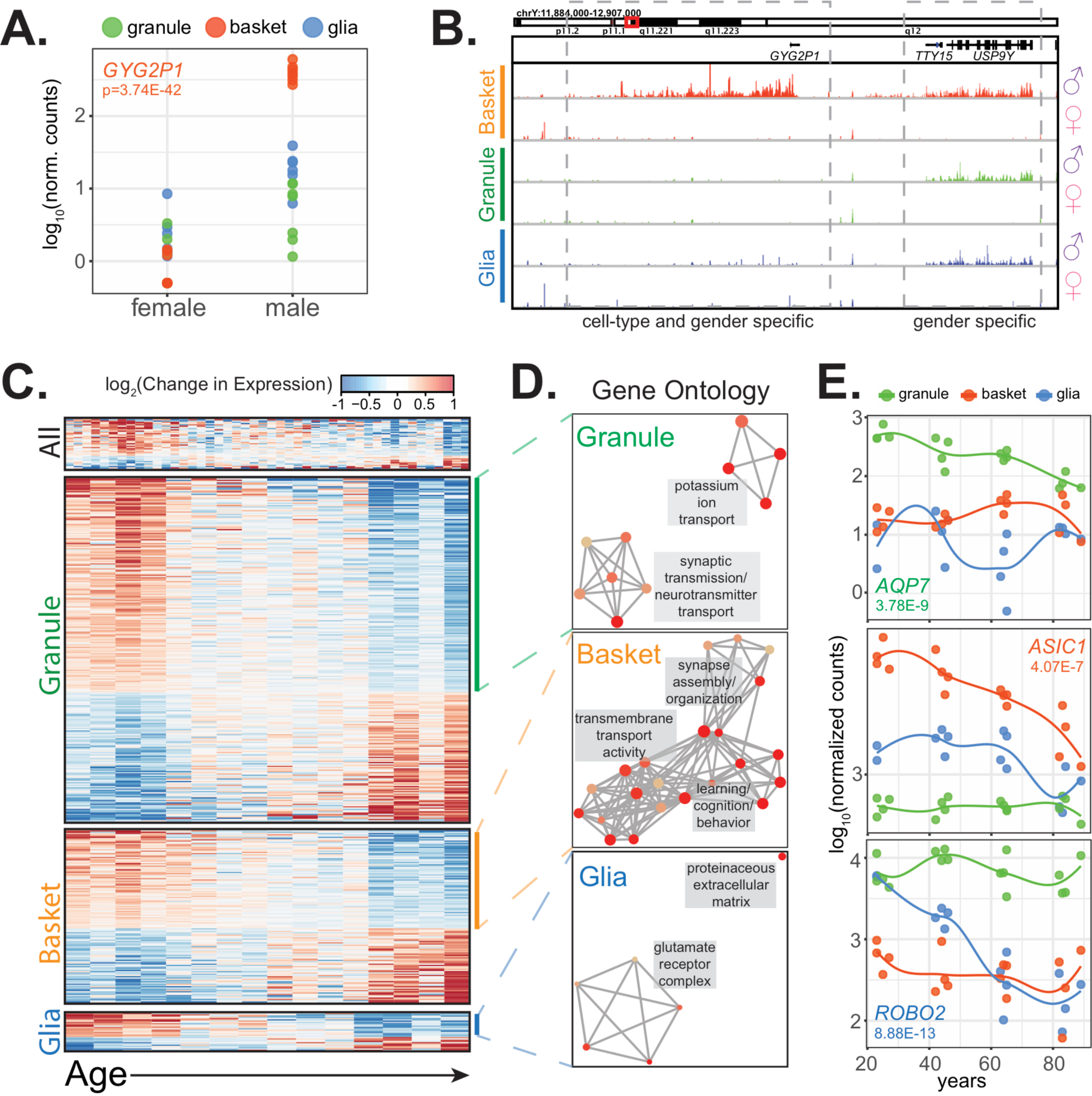
Clinical factors impact gene expression in a cell-type specific manner. **(A,B)** *GYG2P1*: a basket-specific, male-specific gene. **(A)** quantification of *GYG2P1* gene expression in granule, basket, and glial nuclei for female and male samples. **(B)** browser view showing expression of male-specific genes *TTY15* and *USP9Y* and the male-specific and basket-specific gene *GYG2P1*. **(C-E)** Genes that significantly change with age. **(C)** Heatmap showing change in gene expression across all, granule, basket, or glia nuclei. Columns: samples are sorted by age. Rows: genes ranked by change in gene expression over age. **(D)** Enrichment map of significant gene ontology categories for aging down-regulated genes from granule cells, basket cells, and glia. **(E)** Scatterplot showing gene expression across age for three genes that are cell-type specific and down-regulated with age. See also Figure S7.

### Age

A fundamental question in aging research is whether aging occurs uniformly across the brain, or whether some cell types are more susceptible to the effects of aging than others. To address this issue, we examined the effects of age on gene expression in each of the three cerebellar cell types. We used DESeq2 (Love et al., 2014) to find genes that significantly change with age in all samples, or specifically in granule cells, basket cells, or glia. We found 274 significant genes in granule cells, 139 in basket cells, 31 in glia, and 42 that are significant across all samples regardless of cell type (Figure S7A, Table S5). These genes are about equally split between those that increase in expression and those that decrease in expression with age (Figure 6C, S6G). Interestingly, only a few genes significantly change with age in more than one cell type. For example, the 42 genes that are identified as significantly changed with age across all cell types appear heterogeneous across conditions (Figure 6C), and most of these do not reach significance when tested in each cell type individually (Figure S7G). These data suggest that the specific molecular consequences of aging, at least in the cerebellum, are different and depend on the cell type.

Despite differences in the genes impacted by aging in each cell types, it remained possible that the cellular processes altered by these changes are related between cell types. To test this hypothesis, we performed GO analysis on the genes that are up- or down-regulated with age in each cell type. We did not find significant GO categories for any group of aging up-regulated genes, nor did we find significant GO categories for the aging up-downregulated genes from the all cell types group. Interestingly, the genes down-regulated with age in all three individual cell types, despite containing different specific genes, were enriched for synaptic genes (Figure 6D, E). Our finding that genes encoding synaptic components decline in expression with age is consistent with previous findings that axons and dendrites atrophy in the brains of older individuals (Burke and Barnes, 2006; Dickstein et al., 2013; Freeman et al., 2008). However, our data extend these observations to establish that this general process impacts distinct genes in each cell type, as might be expected from the distinct protein compositions of synapses or synapse related functions in each cell types.

### Stress

Transcriptional induction of immediate early genes such as *c-FOS* and *JUNB* is associated with neuronal activation, and can indicate normal behavioral responses or pathological influences (Okuno, 2011; Pérez-Cadahía et al., 2011). Although immediate early response genes associated with these events are often shared, the target genes and pathways induced by these factors reflect the nature of the event experienced by the cell or circuit (Duclot and Kabbaj, 2017). We noticed in our data that *c-FOS* is induced strongly in granule cells obtained from three (WI, VM, and ZH) of the sixteen samples (Figure 7C). To further explore this difference, we performed differential expression analysis to identify additional genes that are significantly up- or down-regulated in these three individuals compared to the others (Figure 7A, Table S5). A very strong response was evident in granule cells from samples WI, VM, and TM, characterized by induction of 224 genes and diminished expression of 10 genes. An overlapping but weaker response that included 136 induced genes was present in glial cells. The induction of these genes in glia is not due to contamination by granule cells as highly expressed markers for granule cells (e.g. *FAT2*) are not present in the glial data. This is interesting because induction *c-FOS* expression by glutamate in glia it is thought to occur via a different mechanism than that induced by increased activity in cultured granule neurons (Edling et al., 2007). Only two differentially expressed genes were evident in the basket cell data.

**Figure 7.**
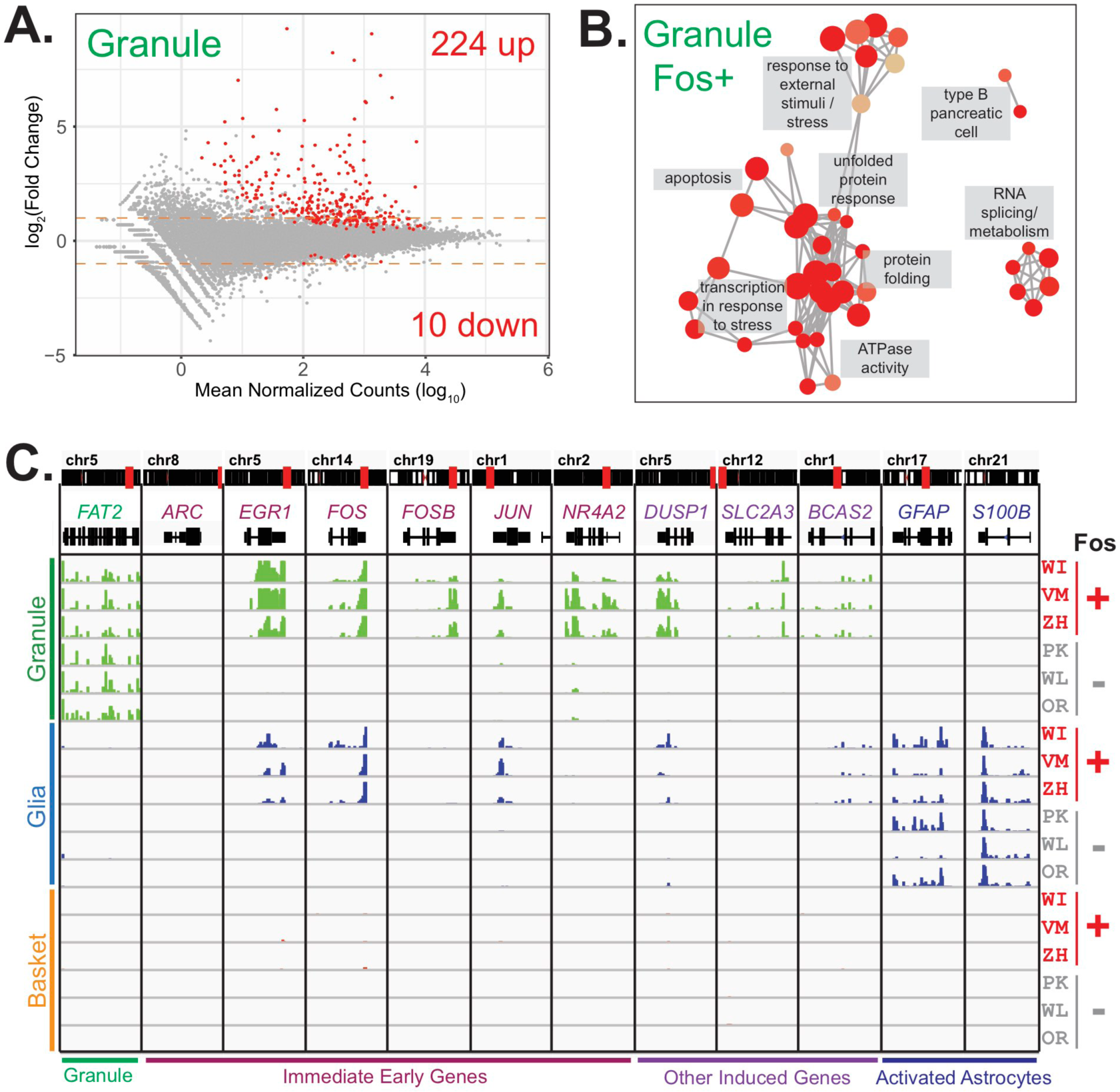
Expression of stress response and immediate early genes in three human samples. **(A)** MA-plot showing differential gene expression analysis of granule cells between three fos+ samples versus the other 13 samples. **(B)** Network diagram of significant enriched GO categories for up-regulated genes in fos+ samples reveals enrichment for genes involved in stress response and protein folding. **(C)** Browser view showing expression of selected genes in three cell types from Fos+ (WI,VM,ZH) and Fos- (PK,WL,OR) human samples that are age and gender matched.

Given the very robust response evident in the granule cell data, we performed GO analysis to determine whether the 224 induced genes in these neurons are enriched for specific functional categories. This analysis revealed categories associated with responses to external stimuli, protein folding/refolding, apoptosis, transcriptional response to stress, ATPase activity, and RNA splicing/metabolism (Figure 7B, Table S6). Analysis of immediate early gene expression indicated that their induction is also selective. For example, induction of *EGR1*, *FOS*, and *FOSB* occurs, but no difference in expression of ARC (a well-characterized immediate early gene) is evident (Figure 7C). Furthermore, although expression of immediate early genes such as *c-FOS* can become induced in response to damage such as ischemic stroke, we found no evidence for induction of glial genes such as GFAP and S100B that mark astrocyte activation in response to stroke and other forms of damage to the brain (Choudhury and Ding, 2016; Ding, 2014; Dirnagl et al., 1999; Kajihara et al., 2001; Pekny and Nilsson, 2005). Finally, examination of the clinical records failed to reveal features (gender, age, cause of death) that distinguished these three samples from the 13 other control donors. Although the cause or causes of these molecular events remain obscure, the discovery of this strong and highly reproducible stress response in select post-mortem human brain samples is consistent with the presence of an occult clinical condition, and indicates that nuclear expression profiling of specific cell types can be used to discover pathways and biomarkers associated with human pathophysiological conditions.

## Discussion

The genes expressed in each cell type define their function, their responses to internal and external cues, and their evolution across species. Here we report a robust strategy for cell type specific expression and epigenetic profiling of human post-mortem brain that incorporates features of several prior protocols for single cell or population based studies of CNS cell types. We demonstrate that cell specific membrane proteins can often be used to sort isolated nuclei, increasingly substantially the candidate antibodies that can be employed for this purpose. We report highly accurate and comprehensive cell-type specific gene expression profiling of neurons and glia from mouse, rat, and human brains. We find that even classically defined, uniform cerebellar cell types differ between species by expression of hundreds of orthologous genes, and that continued expression of human granule cell expressed genes in other regions of the mouse brain commonly occurs. Analysis of samples from sixteen human postmortem brains reveals that the specific molecular consequences of aging differ between cell types, although in each case, expression of genes involved in synapse development and maintenance is diminished. Finally, we report that a robust and cell-type specific molecular pathway indicative of a pathophysiological response occurred in the brains of three of sixteen donors. Taken together, our data provide a resource for comparative analysis of a variety of cell types in rodent and human brains, and they demonstrate that our approach can provide an improved avenue for investigation of molecular events associated with human cell type function and dysfunction.

### Species and cell type specific gene expression in the mammalian brain

Although a consensus definition of cell type has yet to emerge for the mammalian CNS, a recent evolutionary model (Arendt, 2008; Arendt et al., 2016) is helpful for consideration of the data we have generated here. According to this model, the specific characteristics of homologous cell types can vary as long as they remain defined by a distinctive, shared regulatory apparatus. For example, granule cell or astrocyte gene expression profiles can vary between species, or even in individual cells of a type, without losing their cell type identity. This definition can accommodate both functional changes in cell types between species and altered expression of genes within the cell type due to mutations in cis-regulatory sequences. Our data document major differences in the expression of orthologous genes in each cell type between rodent and human brains. Hundreds of these differences are cell-type specific and, in granule neurons, most of these genes remain expressed in other regions of the brain. We believe these data are exciting and provocative because they demonstrate that the fine-tuned biochemistry of homologous human and mouse CNS cell types is likely to differ significantly.

The results presented here contribute two additional insights into mammalian brain evolution. First, while they are consistent with previous published examples of cis-regulatory sequence divergence as a mechanism of evolution (Maricic et al., 2012; Prud'homme et al., 2006; Weyer and Paabo, 2015), the lack of highly enriched GO categories for most of the cell specific events we have documented suggests strongly that regulatory changes are major drivers of phenotypic variation between homologous cell types in mammalian brain. This is consistent with the evolutionary model presented above, since it allows substantial evolution of the functional properties of the cell while maintaining the core regulatory complex (CoRC) that defines a given cell type (Arendt et al., 2016). Second, the expression differences we have documented are large, and we have restricted our analysis to high confidence orthologous genes. Our data, therefore, complement a previous study of progenitor cells that reported substantive changes in the expression of human genes that lack mouse orthologues (Florio et al., 2015). If taken together, these datasets predict that the expressed differences between mouse and human cell types are extensive, cell-type specific, and phenotypically important.

### Molecular phenotyping of human CNS cell types

Recent advances in technologies for human genome sequencing and analysis have led to an astounding increase in our knowledge of the complex genetic causes of human psychiatric and neurological disease (Burguière et al., 2015; Hinz and Geschwind, 2017; Vorstman et al., 2017). In some cases, recreation of the causative mutations in the mouse genome has resulted in experimental models that are sufficiently accurate for investigation of molecular basis of the disorder (Lombardi et al., 2015; Lyst and Bird, 2015; Orr, 2012), and they have led to the discovery of unexpected features of the disease (Guy et al., 2007). In other cases, informative animal models remain elusive (Lavin, 2013), and investigation of molecular mechanisms of disease difficult (Biton et al., 2008; Medina and Avila, 2014). Here we present data for three cerebellar cell types in sixteen human brain samples. All samples were obtained from unaffected subjects that received a neuropathological diagnosis of “normal adult brain”. The correlation coefficients between samples (r=0.98 – 1.0) indicated that the data are extremely reproducible, and that the impact of autolysis time on the expression profile of nuclear RNAs is minor. Initial indications that gender and age impact gene expression differently in CNS cell types are interesting, and they suggest that nuclear profiling is sufficiently accurate for detailed investigation of both sexually dimorphic human behaviors and the aging of specific brain circuitry. An exciting possibility, for example, is that comparative analysis of human brains may provide critical insight into the impacts of aging on cell types that are selectively vulnerable into late-onset human disorders.

One of the most intriguing findings we present is that cerebellar granule cells from three brain samples exhibit robust induction of a set of 224 genes containing many immediate early genes, and that are enriched for GO categories (protein folding/refolding, apoptosis, transcriptional response to stress, response to external stimuli, ATPase activity, etc..) that indicate an acute response to some external influence; yet no shared clinical conditions are apparent for these donors. As stated above, no correlation with autolysis time and no induction of glial markers indicative of stroke or brain damage is evident in our data. Although activity dependent gene expression changes have been characterized extensively in cultured mouse granule cells, the response identified in these three human samples is overlapping but distinctive. While we do not wish to speculate on the cause of this response, our data provide guideposts for experimental investigation of model systems or iPS cells aimed at identification of signals that might have elicited these responses in the human brain.

### Concluding Remarks

The driving force for development of an accurate and efficient non-genetic method for characterization of the molecular properties of defined CNS cell types is to enable direct investigation of human biology. Given the tremendous explosion of studies identifying the genetic causes of human disease and the astounding demonstrations in both animal models and humans that these can include lesions that involve simple changes in gene dosage, exploration of details of cell types in normal and affected brains is imperative. We believe comparative analysis of human cell type specific data with that obtained in experimental systems is an important approach toward advancing our understanding of the large variety of human disorders that impact CNS circuits.

## Author Contributions

Conceptualization, X.X. and N.H.; Methodology, X.X. and N.H.; Software, X.X.; Validation, X.X.; Formal Analysis: X.X. and E.I.S.; Investigation, X.X., E.I.S., A.L., and J.X.; Resources, D.M.; Data Curation, X.X.; Writing - Original Draft, X.X. and N.H.; Writing Review & Editing, X.X. and N.H.; Visualization X.X. and E.I.S.; Supervision N.H; Funding Acquisition X.X. and N.H.

## Acknowledgments

We thank members of the Heintz lab for helpful discussion, suggestions, and improvements. We thank Eric Schmidt, Dakota Blackman and Winrich Freiwald for the rat tissue. We thank Svetlana Mazel, Selamawit Tadesse, Stanka Semova, Songyan Han, and Xiao Li from The Rockefeller University Flow Cytometry Resource Center. We thank Connie Zhao and The Rockefeller University Genomics Resource Center. This work was supported by the HHMI (to N.H.), the NIH/NIDA (grant 1P30 DA035756-01 to N.H.), the Leon Black Family Foundation (to N.H.), the Robertson Therapeutic Development Fund (to N.H. and X.X), and the Estelle G. Kestenbaum Award for Innovative Research in Neurodegenerative Disease. X.X. and N.H. are named as inventors on a patent application filed on this technology, assigned to The Rockefeller University. N.H. is a founder of a startup company using this technology where X.X. is currently an employee. Conduct of the studies presented in this manuscript were not influenced by these relationships.

## METHOD DETAILS

### Nuclei isolation

Nuclei isolation was adapted from the protocol described in previous publications (Kriaucionis and Heintz, 2009; Mellén et al., 2012). For mouse experiments, cortex and cerebella were dissected as described previously. All tissue from C57BL/6J animals were flash frozen using liquid nitrogen. Tissue from bacTRAP animals were freshly dissected and directly used for homogenization. Rat and human tissue were obtained frozen. All frozen tissues were thawed on ice for a minimum of 30 minutes before use.

To isolate nuclei, fresh or thawed tissue were transferred to 5mL of homogenization medium (0.25 M sucrose, 150 mM KCl, 5 mM MgCl2, 20 mM Tricine pH 7.8, 0.15 mM spermine, 0.5 mM spermidine, EDTA-free protease inhibitor cocktail, 1mM DTT, 20U/mL Superase-In RNase inhibitor, 40U/mL RNasin ribonuclease inhibitor). Tissue were homogenized by 30 strokes of loose (A) followed by 30 strokes of tight (B) glass dounce. Homogenate was supplemented with 4.6mL of a 50% iodixanol solution (50% Iodixanol/Optiprep, 150 mM KCl, 5 mM MgCl2, 20 mM Tricine pH 7.8, 0.15 mM spermine, 0.5 mM spermidine, EDTA-free protease inhibitor cocktail, 1mM DTT, 20U/mL Superase-In RNase inhibitor, 40U/mL RNasin ribonuclease inhibitor), and laid on a 27% iodixanol cushion. Nuclei were pelleted by centrifugation 30 min, 10,000 rpm, 4°C in swinging bucket rotor (SW41) in a Beckman Coulter XL-70 ultracentrifuge. For mouse, rat, and human cytoplasmic fractions, 175uL of the top layer was added to 175uL Qiagen Buffer RLT and stored at −80°C. The nuclear pellet was resuspended in homogenization buffer.

### Nuclei labeling and sorting

After nuclei isolation, resuspended nuclei were fixed with 1% formaldehyde for 8 minutes at room temperature, and then quenched with 0.125M glycine for 5 minutes. Nuclei were pelleted at 1000g, 4 minutes, 4°C, and then washed two times with Wash Buffer (PBS, 0.05% TritonX-100, 50ng/mL BSA, 1mM DTT, 10U/uL Superase-In RNase Inhibitor). Nuclei were blocked with Block Buffer (Wash buffer with an additional 50ng/mL BSA) for 30 minutes, incubated with primary antibody for 1 hour, and then washed three times with Wash Buffer with spins in between washes as described above. Nuclei were then incubated in secondary antibody for 30 minutes and washed three times with Wash Buffer. All incubations steps were performed at room temperature. Primary and secondary antibodies were diluted in Block Buffer. Secondary antibodies were purchased from Life Technologies or Jackson Immunoresearch and were used at 1:500 dilution. Secondary antibodies from goat used: mouse Alexa488, rabbit Alexa594. Secondary antibodies from donkey used: mouse Alexa488, goat Alexa488, rabbit Alexa488, mouse Alexa594, rabbit Alexa594.

#### Primary antibodies

| Antigen | Species | Vendor | Cat. # | Dilution |
| --- | --- | --- | --- | --- |
| NeuN | Rabbit | Abcam | ab177487 | 1:250 |
| NeuN | Mouse | Millipore | MAB377 | 1:250 |
| Itpr1 | Mouse | Abcam | ab190239 | 1:500 |
| Sorcs3 | Goat | Thermo | PA5-48023 | 1:250 |
| EEAT1 | Rabbit | Abcam | ab416 | 1:250 |
| Olig2 | Goat | R&D | AF2418 | 1:250 |

Antibodies used for isolating all cell types are below. Unless indicated, NeuN refers to the rabbit antibody.

#### Mouse

Granule: Itpr1, Sorcs3, Olig2, NeuN. Purkinje: Itpr1, NeuN. Basket: Sorcs3, NeuN. Astrocyte: EEAT1, NeuN (mouse). All oligodendroctyes, mature oligodendrocyte, OPC: Olig2, NeuN.

#### Rat

Granule: Itpr1, Sorcs3, Olig2. Purkinje and basket: Itpr1, NeuN. Astrocyte and oligodendrocytes: Sorcs3, NeuN. OPC: Olig2, NeuN.

#### Human

Granule, basket, and glia: Itpr1, NeuN. Astrocyte: Itpr1, Olig2, NeuN. Mature oligodendrocyte, OPC: Olig2, NeuN.

### Flow cytometry

Prior to flow cytometry, nuclei were co-stained with DyeCycle Ruby to 20uM final concentration. Nuclei were analyzed using a BD LSRII (BD Biosciences, San Jose, CA, USA) flow cytometer using the 488nm, 561nm, and 640nm lasers. Nuclei were sorted using a BD FACSAria cell sorter using the 488nm, 561nm, and 635/640nm lasers. All samples were first gated using DyeCycle Ruby to determine singlets. Analysis was performed using FACSDiva (BD) or FlowJo software. Qiagen Buffer RLT (unfixed samples) or Qiagen Buffer PKD (fixed samples) were added to the sorted nuclei and the samples were stored at −80°C. Information about percentages for all gated populations can be found in Table S1.

### RNA purification, library construction, and sequencing

RNA from cytoplasmic fractions or fresh nuclei were purified using the Qiagen RNeasy Micro kit with on-column DNase digestion. RNA from fixed nuclei were purified using the Qiagen RNeasy FFPE kit with the following modifications – after Proteinase K digestion, RNA was spun at max speed for 15 minutes. The supernatant was removed and incubated at 65°C for 30 minutes, 70°C for 30 minutes, and 80°C for 15 minutes. An on-column DNase digestion was performed in place of the in-solution DNAase digestion. RNA quantity was determined using the Qubit RNA HS Assay kit and RNA quality was determined using Agilent 2100 Bioanalyzer with RNA 6000 Pico chips. Purified RNA was converted to cDNA and amplified using the Nugen Ovation RNA-seq System V2. cDNA was fragmented to an average size of 250bp using a Covaris C2 sonicator with the following parameters: intensity 5, duty cycle 10%, cycles per burst 200, treatment time 120 seconds. Libraries were prepared using the Illumina TruSeq DNA LT Library Prepartion Kit or the NEBNext Ultra DNA Library Prep Kit for Illumina with NEBNext Multiplex Oligos for Illumina. The quality of the libraries was assessed using the Agilent 2200 TapeStation system with D1000 High Sensitivity ScreenTapes. Libraries were sequenced at The Rockefeller University Genomics Resource Center on the Illumina HiSeq 2500 machine to obtain 50bp single-end reads or on the Illumina NextSeq 500 to obtain 75bp paired-end reads. A summary of all RNA-seq datasets can be found in Table S1. All datasets have been deposited in GEO: accession-numbers-pending.

### ATAC-seq library construction and sequencing

ATAC-seq libraries were prepared as described (Buenrostro et al., 2013; 2015), with minor modifications. Nuclei were extracted as described above and incubated for 10 minutes in lysis buffer (10 mM Tris pH 7.5, 10 mM NaCl, 3mM MgCl2, 0.1% NP-40). Nuclei were then resuspended in 50 ul 1x TD Buffer containing 2.5 ul Tn5 enzyme from Nextera kit (Illumina). Transposition reactions were incubated for 30 min at 37°C and purified with QiaQuick MinElute columns (Qiagen). Purified DNA was amplified by PCR using Q5 High Fidelity Polymerase (NEB) for 12 cycles with barcoded primers, as described (Buenrostro et al., 2013). Amplified libraries were purified and size selected with AMPure XP beads (Beckman Coulter). Libraries were sequences on Illumina HiSeq 2500 to yield 50 bp paired-end reads and on Illumina NextSeq 500 to yield 75 bp paired-end reads.

### Immunofluorescence

Mice were deeply anesthetized and transcardially perfused with phosphate buffered saline (PBS) followed by 4% formaldehyde (w/v) in PBS. Brains were dissected and postfixed overnight at 4°C, cryoprotected in 30% sucrose in PBS, OCT TissueTeck embedded, and cut with a Leica CM3050 S cryostat into 20um sections that were directly mounted on slides. Slides were stored at −20°C. Antigen retrieval was performed by immersing slides into sodium citrate buffer (10mM sodium citrate, 0.05% Tween 20, pH 6.0) at 95-100°C and simmering for 10 minutes in the microwave. The slides were cooled to room temperature and then washed with PBS. Immunofluorescence was performed by blocking for 30 minutes in IF Block Buffer (3% BSA in PBS with 0.1% TritonX-100), incubated with primary antibody overnight, washed 3 times with PBS, incubated with secondary antibody for one hour, washed with PBS, stained with DAPI solution (1ug/mL in PBS) for 15 minutes, washed two times with PBS, and coverslipped with Prolong Diamond mounting media. For some samples, Tyramide signal amplification was performed as follows: slides were incubated with secondary antibody conjugated to horseradish peroxidase for 1 hour, washed 3 times with PBS, incubated with Cy3 in Amplification Buffer from the TSA Cyanine 3 detection kit for 10 minutes, washed with PBS, and DAPI stained as above. All steps were performed at room temperature. Slides were imaged on a Zeiss LSM700 confocal microscope using the same acquisition settings for mouse and human slides. Brightness and contrast adjustments were made in ImageJ post-acquistion with the same adjustments applied to mouse and human images. In addition to the antibodies listed above for nuclei staining, the following primary antibodies were used for immunostaining:

#### Primary antibodies

| Antigen | Species | Vendor | Cat. # | Dilution |
| --- | --- | --- | --- | --- |
| GFAP | Goat | Abcam | ab53554 | 1:500 |
| Mog | Goat | Thermo | PA5-47319 | 1:500 |
| Mag | Rabbit | Cell Signaling | 9043S | 1:500 |
| Pde1a | Goat | Acris | TA311317 | 1:200 |
| Pde1c | Rabbit | Santa Cruz | sc-376474 | 1:100 |

### Software used for analysis

Data processing steps made use of Linux tools in addition to custom perl scripts. In addition to standard Linux commands (e.g. awk, cut, sort, uniq), we used the following packages: trim_galore (v0.4.1) for adapter trimming of ATAC-seq samples, STAR (v2.4.2a) and Bowtie2 (v2.1.0) for read alignment, SAMtools (v0.1.19-44428cd) for indexing and removal of duplicates, igvtools (v2.3.32) for generating files for browser visualization, deepTools (v2.0) for analysis of ATAC-seq samples, and featureCounts (v1.5.2) for read summerization. Data analysis was performed using R Studio (v 1.0.143, R: v3.4.0). In addition to base R (for data wrangling, Pearson correlations, hierarchical clustering, etc.) and custom R functions, we made extensive use of the following packages: tidyverse (v1.1.1, which contains ggplot2, reshape2, etc.) for data wrangling and visualization, DESeq2 (v1.16.1) for raw count normalization, differential expression analysis, and principle components analysis, gplots (v3.0.1) and pheatmap (v1.0.8) for heatmaps, RColorBrewer (v1.1-2) for color palettes, and clusterProfiler (v3.4.3) for GO analysis.

### ATAC-seq read mapping, visualization, and analysis

Reads were processed with trim_galore with parameters “--stringency 3 --fastqc --paired”. Trimmed reads were mapped to mm10/hg38/rn6 using bowtie2 (Langmead and Salzberg, 2012) with default parameters. Duplicates were removed using samtools (Li et al., 2009). Reads were normalized to 1x depth (reads per genome coverage, RPGC) using deepTools (Ramírez et al., 2016) bamCompare module, ignoring chrX, chrY, chrM and filtering reads for minimum mapping quality of 30. Metagene and heatmap profiles were generated using deepTools modules computeMatrix, plotProfile and plotHeatmap.

### RNA-seq read mapping, visualization, and read summerization

RNA-seq reads were aligned using STAR (Dobin et al., 2013) and genome assemblies from ENSEMBL. In addition to default STAR parameters, we used the following parameters for single end (--outFilterMismatchNmax 999 --outFilterScoreMinOverLread 0 --outFilterMatchNminOverLread 0 --outFilterMatchNmin 35 --outFilterMismatchNoverLmax 0.05) and paired-end data (--outFilterMismatchNmax 999 --alignMatesGapMax 1000000 -- outFilterScoreMinOverLread 0 --outFilterMatchNminOverLread 0 --outFilterMatchNmin 60 -- outFilterMismatchNoverLmax 0.05). Aligned bam files were converted to tdf format for visualization using igvtools. Raw counts were generated using featureCounts (Liao et al., 2014). Refseq or ENSEMBL gene model annotations for whole genes were downloaded using the UCSC Table Browser tool. While ENSEMBL annotates more genes than Refseq (Zhao and Zhang, 2015), we found that visually, our mouse and human RNA-seq data better matches Refseq than ENSEMBL gene models. Because of this, for within species comparisons for mouse and human, we used Refseq annotations. Rat gene models are incompletely annotated by both Refseq and ENSEMBL, but as fewer genes are missing in the ENSEMBL annotation, we chose this annotation for within species comparisons. For comparative analysis across species, we used the ENSEMBL annotation for all species in order to match the list of orthologous transcripts that we obtained from ENSEMBL. The following parameters were used in addition in addition to the default in featureCounts: for paired-end data, fragments are counted instead of reads (-p), chimeric fragments are not counted (-C). For comparative analysis across species, reads are allowed to be assigned to more than one matched meta-feature (-O) in order to avoid problems when genes overlap in one species but not in another. For within-species comparisons, reads that map to more than one matched meta-feature are not counted (default).

#### Genome/gene annotations

| Purpose | Program | Species | Annotations | Source | Genome |
| --- | --- | --- | --- | --- | --- |
| Genome alignment | STAR/Bowtie2 | Mouse | ENSEMBL | ENSEMBL | mm10 |
| Genome alignment | STAR/Bowtie2 | Rat | ENSEMBL | ENSEMBL | rn6 |
| Genome alignment | STAR/Bowtie2 | Human | ENSEMBL | ENSEMBL | hg38 |
| Read quantification | featureCounts | Mouse | Refseq | UCSC | mm10 |
| Read quantification | featureCounts | Rat | ENSEMBL | UCSC | rn6 |
| Read quantification | featureCounts | Human | Refseq | UCSC | hg38 |
| Ortholog read quantification | featureCounts | Mouse | ENSEMBL | ENSEMBL | mm10 |
| Ortholog read quantification | featureCounts | Rat | ENSEMBL | ENSEMBL | rn6 |
| Ortholog read quantification | featureCounts | Human | ENSEMBL | ENSEMBL | hg38 |

### Comparison with single nuclei RNA-seq (sNuc-Seq) data

Raw sequence files were downloaded from GEO for 17 astrocyte, 8 oligodendrocyte, and 7 OPC single nuclei mouse RNA-seq (sNuc-Seq) datasets from (Habib et al., 2016). To directly compare these datasets to our bulk nuclear profiles for the same cell types, downloaded sequences were aligned to the mm10 genome and raw counts against mouse Refseq whole gene annotation were generated as described above. Log transformed transcripts per million (TPMs) were calculated for both sNuc-Seq and our nuclear profiles as described by (Habib et al., 2016) except that lengths for the whole gene instead of transcripts (exons only) were used for normalization. To explore the effect of sequencing depth on the number of genes detected, the number of mapped reads for single nuclei data were extracted using samtools flagstat. We next used samtools view to randomly downsample our nuclear data to 50, 100, 200, 300, 400, 500, 600, 800, 1,000, and 1,200 thousand reads. We regenerated gene counts using the downsampled data and calculated TPMs. Using the same threshold as (Habib et al., 2016), we considered a gene to be expressed if it has a log_2_(TPM+1) > 1.1.

### Generation of ortholog annotations

Because we were interested in differences in highly conserved cell types, we only wanted to study differences in high confidence orthologs. Schematics for generating annotations for mouse-rat-human orthologs, and mouse-human orthologs are shown in Figures S3A and S4A respectively. In both cases, we first downloaded a list of orthologous transcripts from the ENSEMBL BioMart (release 88). First, we removed any annotation that contained a low confidence pair (ENSEMBL orthology confidence = 0). Next, we removed duplicate annotations as well as any orthologs that were not strictly 1:1, which we defined as transcript that is annotated to be orthologous to more than one transcript in another species. To avoid any differential expression simply due to changes in gene length, we decided to filter out any genes that change in length by more than two-fold across any species. Additionally, as we were not interested in examining small RNAs, we also filtered out genes smaller than 1kb in genome length (including untranslated regions). Finally, because we were interested in gene level rather than transcript level differences, we used the annotation for the longest transcript as the gene annotation. Because the rat genome is incompletely annotated, we ended up with almost 3,000 more genes that are orthologous between mouse and human, and those that are orthologous between mouse, human, and rat.

### Gene count normalization, differential expression analysis, and principle components analysis

Raw gene count tables were loaded into R and differential expression analysis was performed using DESeq2 (Love et al., 2014). For age and autolysis time, numerical covariates were used for modeling. Statistically significant genes were defined as having an adjusted p-value of less than 0.01. Fold change cutoffs were not used unless otherwise noted. The rlog function was used to generate normalized counts for downstream analysis (e.g. heatmaps, clustering). The plotPCA function was used to perform principle components analysis. We did not use any methods to remove batch effects as we always observed clustering based on expected biology, and never by batch. For example, we observe clustering by cell type between EGFP+ and antibody sorted nuclei despite numerous differences: date of preparation, tissue state (fresh vs. flash frozen and thawed), nuclei state (unfixed vs. fixed), nuclei processing (none vs. antibody stained), library prep kits (Illumina vs. NEB), sequencer (HiSeq 2500 vs NextSeq 500), and read type (50bp single end vs. 75bp paired end).

### Hierarchical clustering

Hierarchical clustering was performed using normalized gene counts produced by the rlog function from DESeq2 (Love et al., 2014). Unless specified, the top 250 most variable genes across samples, as computed by the rowVars function, were used for clustering. We use the most variable genes instead of all genes for clustering because in our experience, for comparisons of related cell types such as neuroglia, the most variable genes are the most informative for defining the different functions between cell types. Additionally, our finding that gene expression between different cell types is quite correlated (r=0.91-0.94 for mouse cerebellar cell types, Figure 1F) suggests that clustering using all genes might dilute the strong but relatively few gene expression differences between cell types. Default parameters (Euclidean distance and complete linkage) were used with the dist and hclust functions to produce clusters.

### Specificity index algorithm

The specificity index (SI) algorithm was previously developed in the lab to find genes that are specific in a dataset of interest as compared to other datasets (Dougherty et al., 2010). Because the previous algorithm was designed for microarray data, we decided to adapt it to accept RNA-seq data as input. We have also added a feature that incorporates information from biological replicates for calculations. In our implementation, fpkms are calculated from raw counts to avoid biases toward long genes, and genes less than 1kb are excluded to avoid biases from extremely short genes. Next, fpkm values are logged and log_10_(fpkm) values are bottomed out at −2 to avoid biases from long, non-expressed genes. To take into account sample replicates, we calculate the mean and standard deviation for all genes within a cell type. SI values are calculated as described previously, but instead of using mean gene expression values as input, we randomly sample from a normal distribution with the mean and standard deviation of gene expression. This process is performed 1000 times and final ranks are averaged across all iterations. R code containing this algorithm can be found at: bitbucket-link. For analysis of the distribution of highly specific genes in the mouse cerebellum, images for each gene from the Allen Mouse Brain Atlas (Lein et al., 2007) were examined. Genes that were not in the Atlas at the time of analysis, or genes that showed no staining in any region of the brain were excluded from analysis. The percentage of genes showing proper distribution for the cell type analyzed were calculated.

### GO analysis

Significant differentially expressed genes were split into up- and down-regulated (or mouse and human enriched) genes and gene ontology enrichment analysis for each group was performed using clusterProfiler (Yu et al., 2012). Significant ontologies were defined as having a q-value of less than 0.01 (default). Table S5 contains all GO analysis results, including significant ontologies from any of the three domains (biological process, molecular function, cellular component). For simplicity of visualization, only biological process ontologies were used, except for in Figure 6D lowest panel as only one biological process ontology was significant while six cellular component ontologies were significant. Additionally, when many ontologies were found, the simplify function was used to remove redundant terms for visualization. Network based enrichment maps were generated using the enrichMap function. To simplify visualization of the enrichment maps, we summarized groups of related ontologies into one label.

## SUPPLEMENTAL FIGURE LEGENDS

**Figure S1.**
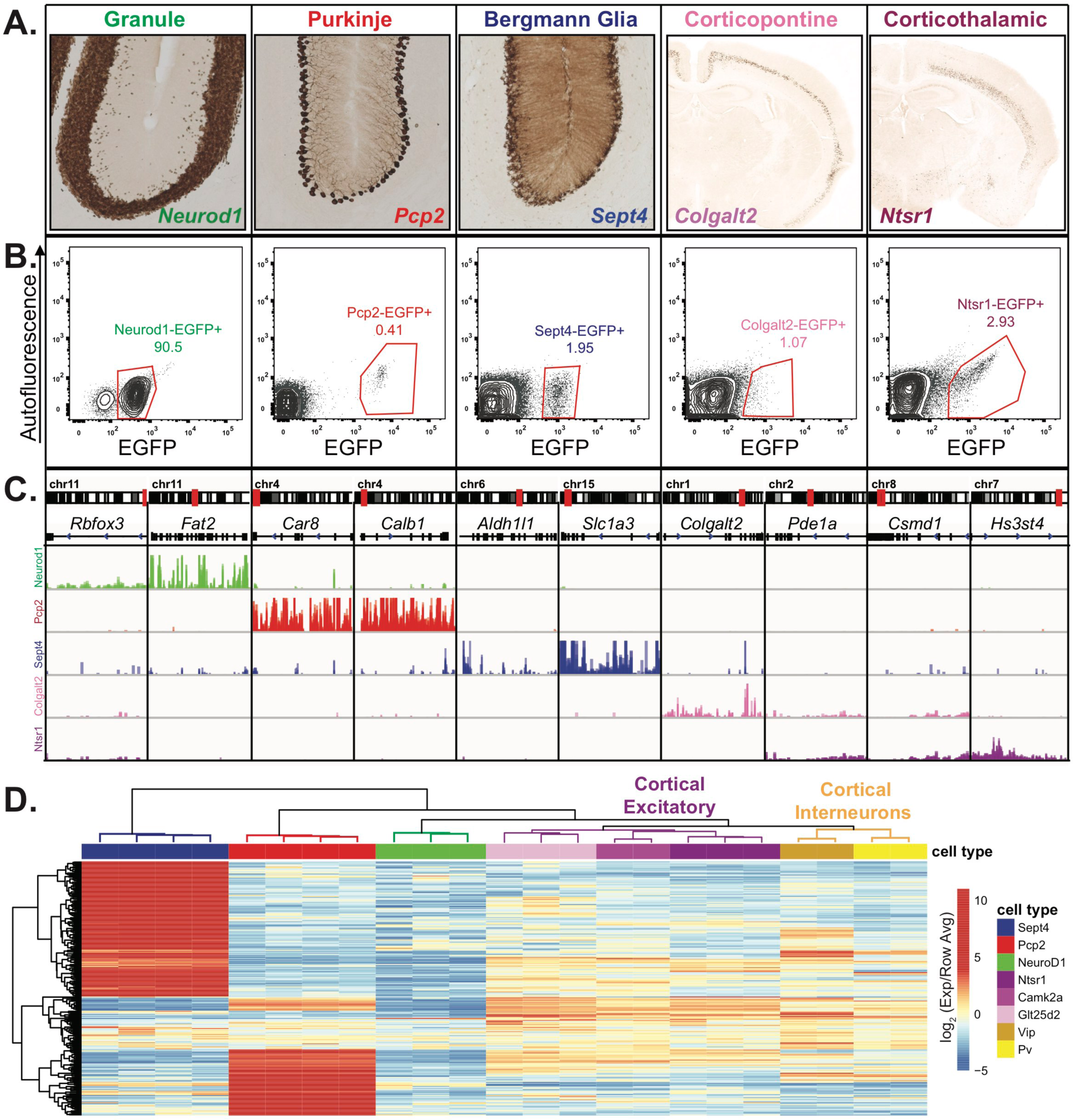
Nuclear RNA profiles can specify cell-type identity, related to Figure 1. **(A)** EGFP-L10a expression from five bacTRAP animals. *Neurod1*, *Pcp2*, and *Sept4* drive expression in granule cells, Purkinje cells, and Bergmann glia of the cerebellum. *Colgalt2* and *Ntsr1* drive expression in corticopontine and corticothalamic pyramidal cells of the cortex. All images are from the GENSAT Project. **(B)** Fluorescence activated sorting for EGFP+ nuclei from each of the five bacTRAP lines. The percentage of GFP+ nuclei is indicated. **(C)** Browser view showing nuclear expression of genes that are specific to each of the five cell types, including genes that are shared between two cell types (*Pde1a* and *Csmd1* are expressed in both cortical pyramidal cell types). **(D)** Heatmap showing expression of the 250 most variable genes across eight cell types. In addition to the five cell types described in **(A-C)**, nuclear expression from three cortical cell types - excitatory neurons, PV interneurons, and VIP interneurons – from (Mo et al., 2015) are shown. Expression is normalized to the average expression across all samples for each gene. Hierarchical clustering is performed on both samples and genes.

**Figure S2.**
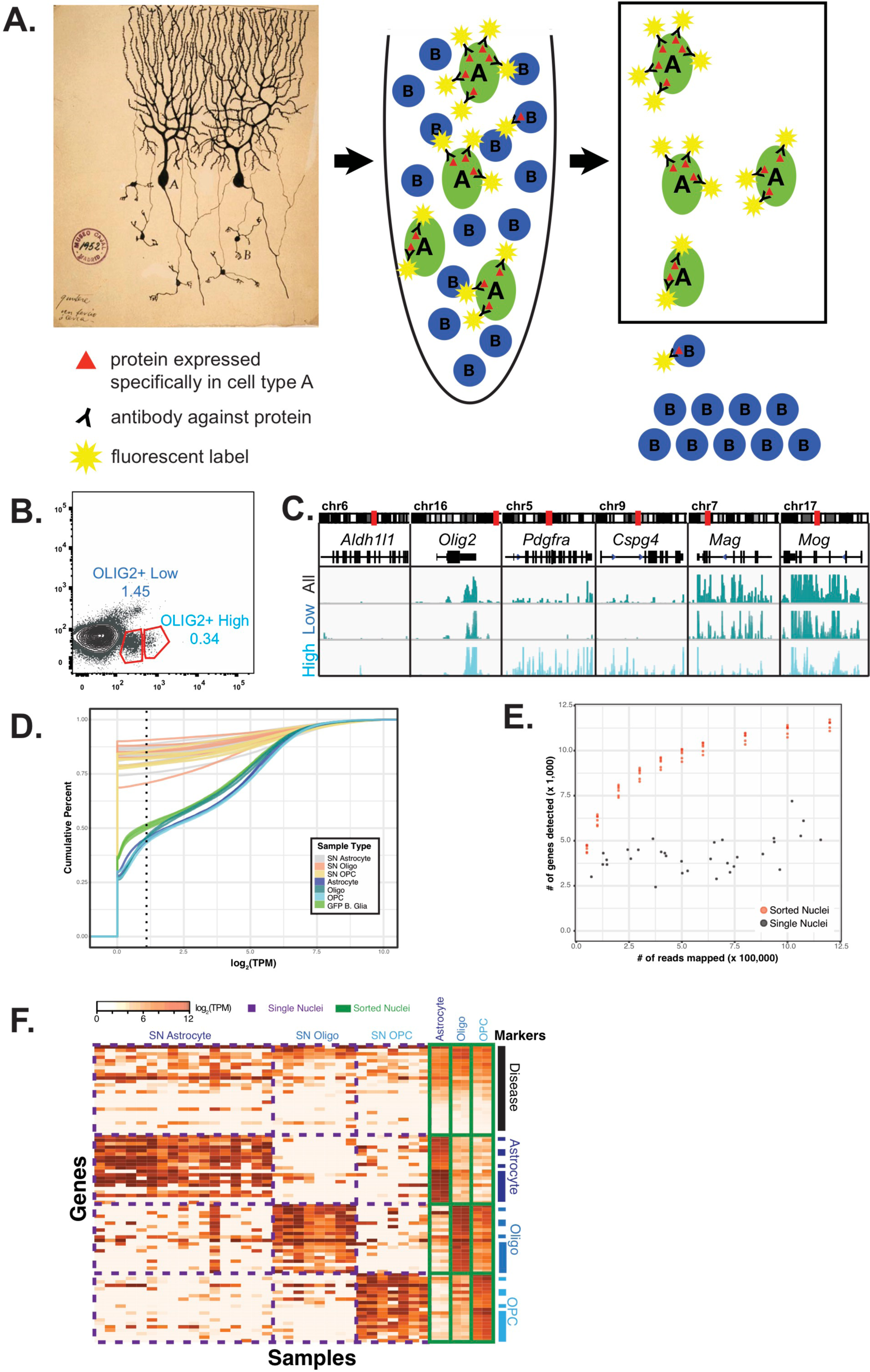
Overview of nuclei labeling and sorting strategy and comparison to single nuclei sequencing, related to Figure 1. **(A)** Schematic showing the strategy for cell-type specific nuclei purification and gene expression profiling. Starting from whole tissue containing heterogeneous cell populations intermingled together, nuclei are isolated. Nuclei from a cell type of interest are fluorescently labeled using antibodies against proteins found specifically in that cell type, and then separated using fluorescent activated nuclear sorting. RNA isolated from these nuclei is used for genome-wide expression profiling by RNA-seq. **(B)** The OLIG2+ population can be separated into two populations. **(C)** Expression profiling of the two populations by RNA-seq reveals that OLIG2+ low nuclei come from mature oligodendrocytes and OLIG2+ high nuclei come from both oligodendrocyte precursor cells (OPCs) and mature oligodendrocytes. Browser view shows gene expression from OLIG2+ low (Mature) and OLIG2+ high (OPC + Mature) populations. Markers: *Aldh1l1* (astrocytes), *Olig2* (all oligodendrocytes), *Pdgra* and *Cspg4* (OPCs), *Mag* and *Mog* (mature oligodendrocytes). (D-E) Comparative analysis of mouse cerebellar glial nuclear RNA-seq data from this study compared to mouse hippocampal single nuclei sequencing (sNuc-Seq) data from (Habib et al. 2016). (D) Cumulative distribution function plot of normalized read counts per gene in log transcripts per million (log_2_(TPM)) for sNuc-Seq (grey/orange/tan) and antibody sorted nuclear RNA-seq datasets. Also shown are 4 replicates of Bergmann glia nuclear RNA-seq datasets obtained from GFP+ sorted nuclei from Sept4-EGFP-L10a BacTRAP mice (green). The dotted line at 1.1 log_2_(TPM) is the threshold used by Habib et al. for defining expressed genes. **(E)** Scatterplot showing number of reads sequenced versus number of genes detected for sNuc-Seq (grey) and down-sampled nuclear RNA-seq datasets (orange). For each sNuc-Seq dataset, the number of aligned reads were extracted using samtools. For each nuclear RNA-seq dataset, samtools was used to randomly sample 50, 100, 200, 300, 400, 500, 600, 800, 1,000, and 1,200 thousand reads. The number of genes detected was defined as the number of genes with a log_2_(TPM) of greater than 1.1. **(F)** Heatmap showing normalized gene expression in sNuc-Seq (purple dashed boxes) and nuclear RNA-seq datasets (green boxes) for neurological disease, astrocyte, oligodendrocyte, and OPC marker genes. Disease genes were taken from a review of Alzheimer’s disease genes (Karch and Goate 2015). Astrocyte, oligodendrocyte, and OPC marker genes were derived from Table S1 from Habib et al. or from Table S2 from this paper. The top ten genes from each group were used: highest by TPM value for the Habib et al. genes or lowest by Specificity Index for nuclear RNA-seq data.

**Figure S3.**
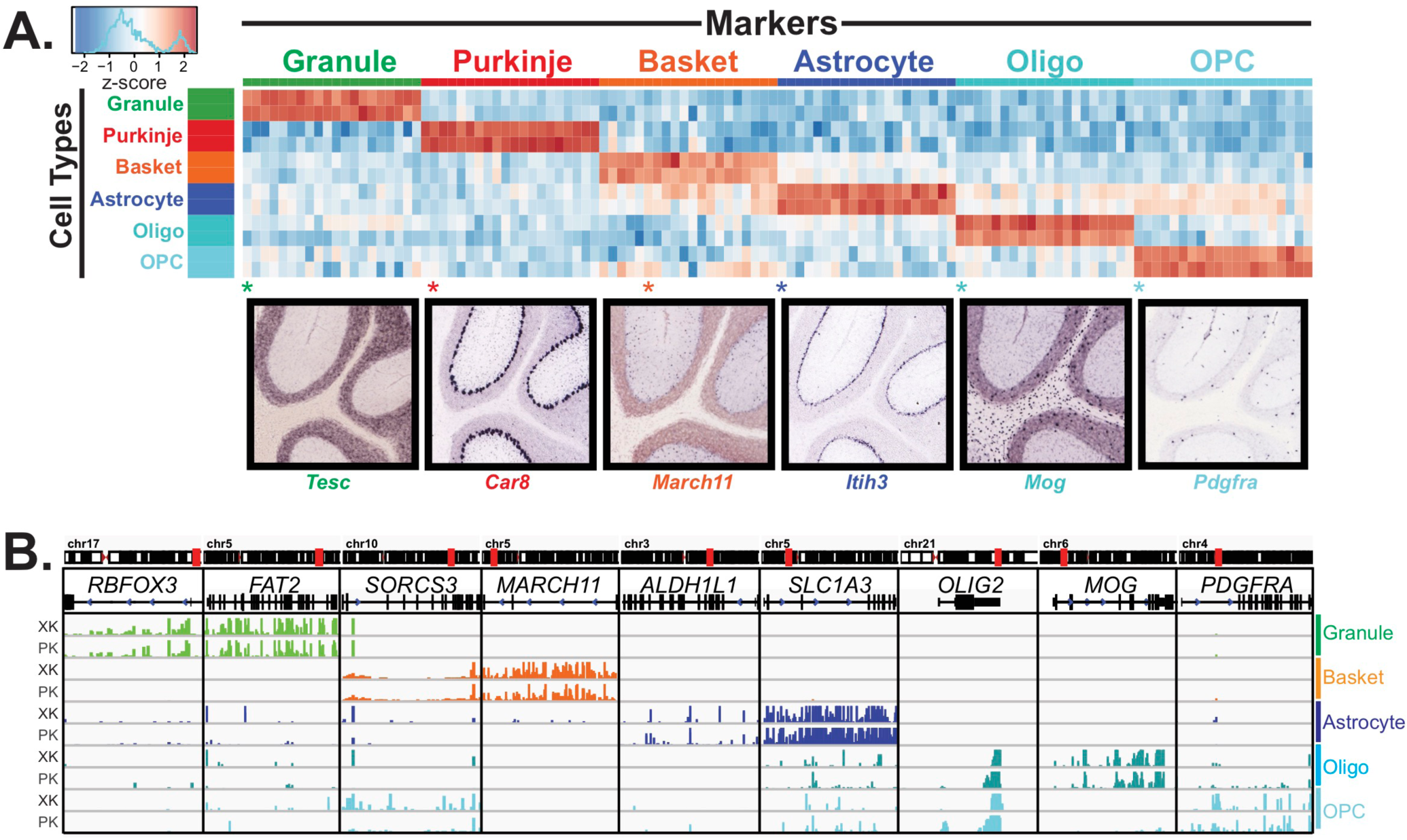
Analysis of cell type specific gene expression profiles generated from rat and human cerebella identifies known mouse marker genes for each cell type. **(A)**Heatmap showing the 20 most specific genes for each rat cell type as identified by the Specificity Index algorithm. Rows: cell type samples; columns: genes. Color represents the z-score of gene expression compared to all samples. Lower panels show examples of gene expression from the Allen Mouse Brain Atlas for highly specific genes identified by analysis of the rat data. Distribution of astrocyte-specific gene *Itih3* is characteristic of both Bergmann glia, a type of astrocyte, and non-Bergmann glia astrocytes of the cerebellum. The percentage of top 20 rat SI genes evident in Allen Mouse Brain Atlas mouse ISH database that show expected distribution for each cell type: granule 93%, Purkinje 100%, basket 85%, astrocyte 93%, oligodendrocyte 100%, OPC 80%. **(B)** Browser view showing gene expression of cell-type specific markers across human samples from five cell types.

**Figure S4.**
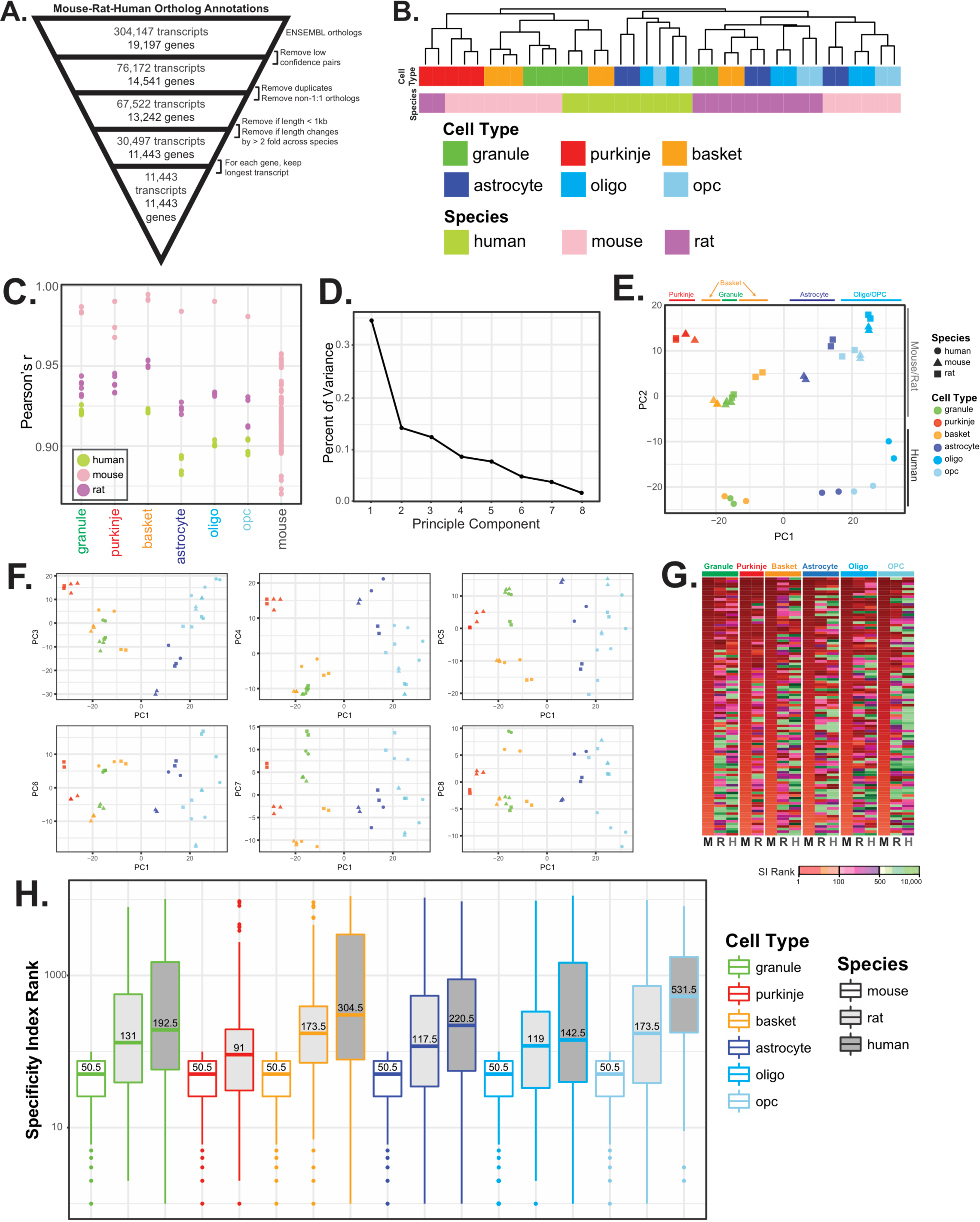
Comparative analysis of gene expression in specific cell types of mouse, rat, and human, related to Figure 3. **(A)** Schematic for defining ortholog annotation for mouse, rat, and human genes. Ortholog annotation across species were downloaded from ENSEMBL, filtered to include only high confidence pairs, 1:1 orthologs, genes greater than 1kb in total length, and genes that change by more than 2-fold in length across species. For each gene, the longest orthologous transcript was used for annotation. **(B)** Hierarchical clustering of mouse, rat, and human cell-type specific datasets based on expression of all genes instead of the top 250 most variable genes reveals clustering primarily by species. **(C)** Dot plot showing pairwise Pearson’s correlation coefficient for each sample, broken down by cell type. Last column shows the pairwise Pearson’s correlation coefficient between all cell types in mouse. **(D-F)** Principle components analysis of all datasets. **(D)** Plot showing the percent of variance explained by each of the first eight principle components. **(E)** Scatterplot showing values of the first two principle components (PC1, PC2) for all samples. PC1 separates samples based on cell type while PC2 separates mouse and rat from human samples. **(F)** Scatterplots showing the first principle component versus components three through eight. **(G,H)** Specificity index (SI) analysis of cell-type specific genes in mouse, rat, and human. **(G)** Heatmap showing SI calculated ranks. Shown are the top 100 mouse SI genes for each cell type. Colors indicate rank position of these 100 genes in mouse, rat, and human. **(H)** Boxplot representation of SI ranks from **(G)**.

**Figure S5.**
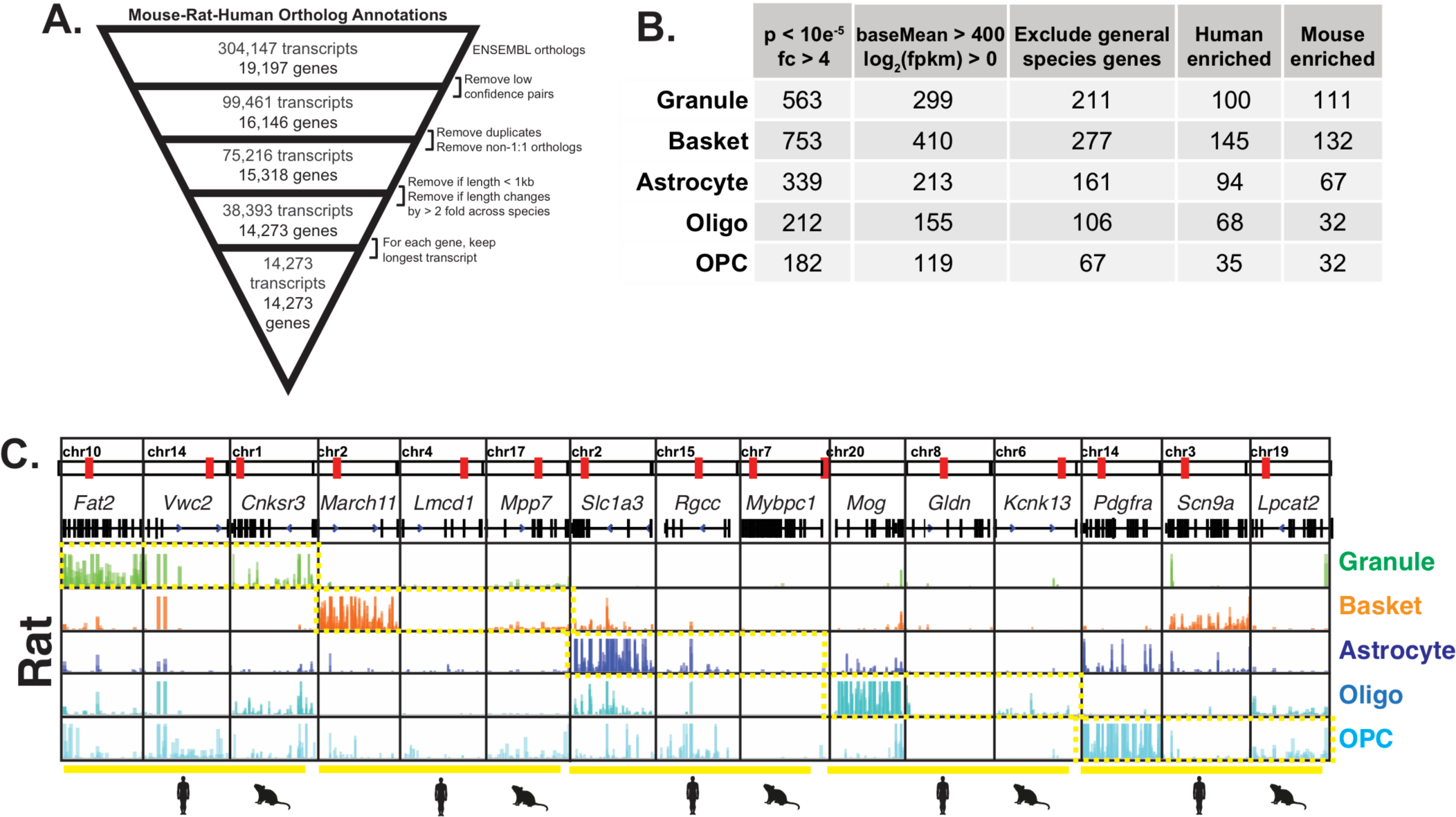
Detailed comparative analysis of gene expression in five cerebellar cell types in mouse and human, related to Figure 4. **(A)** Schematic for defining ortholog annotations for mouse and human genes. Ortholog annotation across species were downloaded from ENSEMBL, filtered to include only high confidence pairs, 1:1 orthologs, genes greater than 1kb in total length, and genes that change by less than 2-fold in length across species. For each gene, the longest orthologous transcript was used for annotation. **(B)** Table Showing for each cell type, the number of genes that are differentially expressed (adjusted p < 10e-5, fold change > 4) between mouse and human, the number that are left after filtering for expression levels (baseMean > 400, log_2_(fpkm) > 0), and the number that are left after filtering for genes that are differentially expressed across all cell types between mouse and human. This number is then broken down into human and mouse enriched genes. **(C)** Browser views showing expression in rat for marker, human-enriched, and mouse-enriched genes.

**Figure S6.**
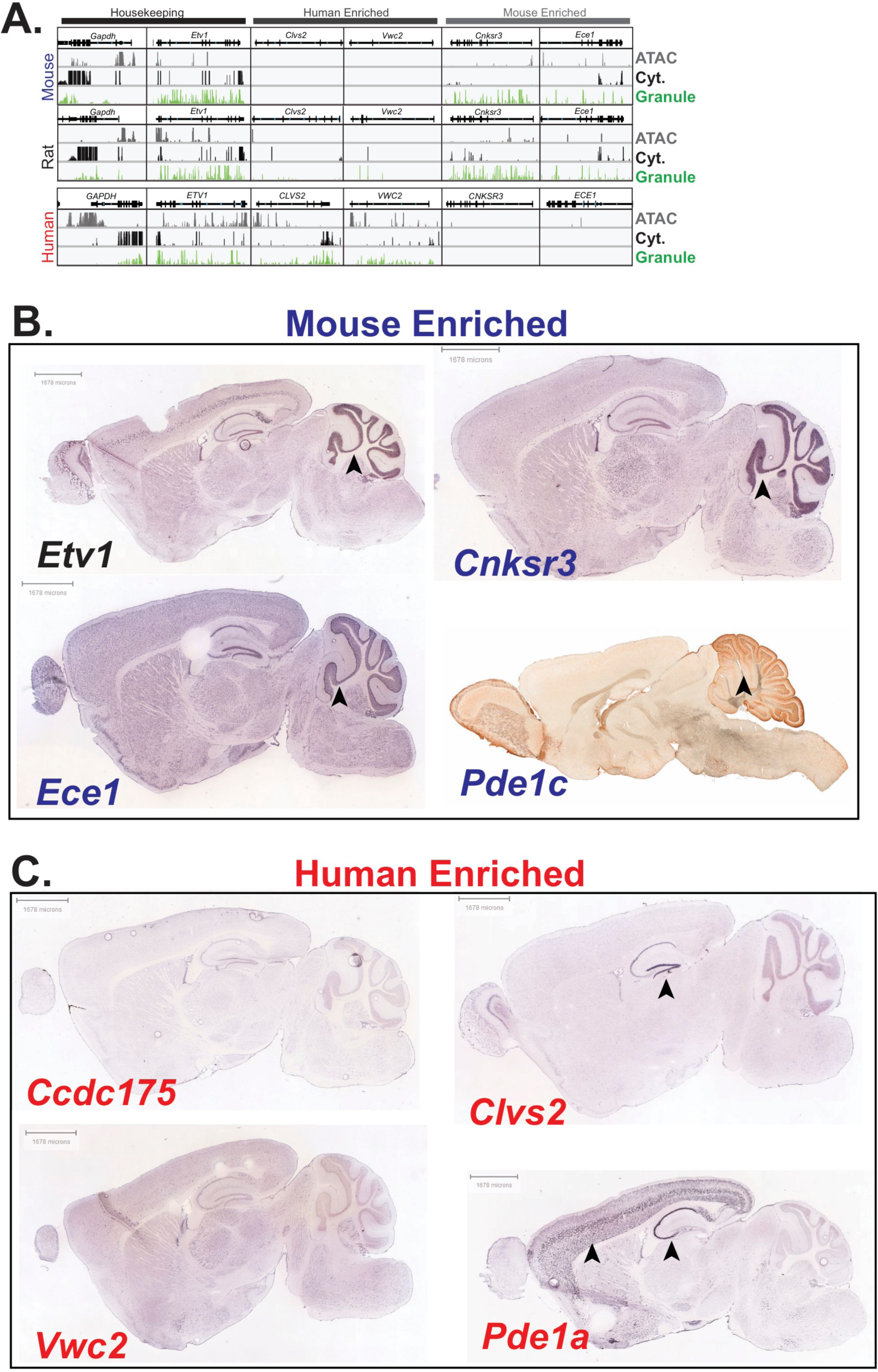
Expression of mouse- and human-enriched granule cell genes in the mouse brain. **(A)** Browser view showing examples of housekeeping, human-enriched, and mouse-enriched genes. Tracks: nuclear RNA levels from granule cells (green), cytoplasmic RNA levels from total cerebellum (black), and ATAC-seq DNA accessbility from total cerebellum (grey) – for mouse, rat, and human. **(B, C)** All images are from the Allen Mouse Brain Atlas, unless noted. **(B)** Expression of the granule-specific marker Etv1 and mouse-enriched genes *Cnksr3*, *Ece1*, and *Pde1c* in the granule layer of the cerebellum (indicated by arrow). *Ece1* expression is also observed in the Purkinje layer of the cerebellum. *Pde1c* expression image is from the GENSAT Project. **(C)** Expression of the human enriched genes *Ccdc175*, *Clvs2*, *Vwc2*, and *Pde1a* in the mouse brain. No staining was detected in any region for *Ccdc175* and *Vwc2*. Arrows point to strong staining of *Clvs2* in the dentate gyrus and of *Pde1a* in layers 5 and 6 of the cortex and CA1-3 of the hippocampus.

**Figure S7.**
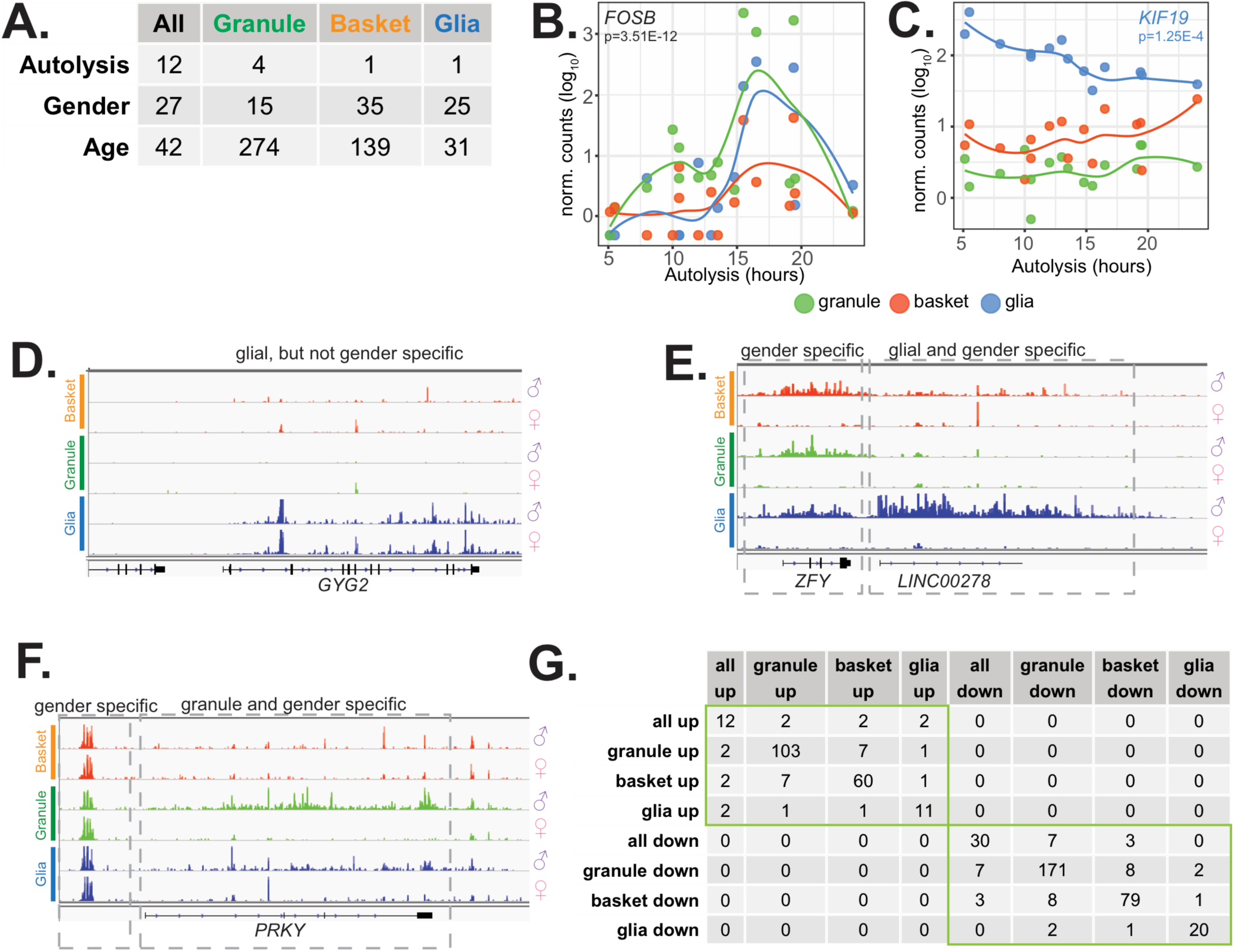
Analysis of clinical factors that contribute to gene expression variability across individuals, related to Figure 6. **(A)** Table Showing the number of differentially expressed genes (adjusted p < 0.01, baseMean > 50) for all cell types or each cell type individually when samples are stratified by clinical factors. For age and autolysis time, numerical covariates were used for differential expression analysis. Gender indicates male versus female comparison. **(B,C)** Examples of genes that vary in expression depending on autolysis time. **(B)** FOSB is elevated in granule and basket cells in a few samples with intermediate autolysis times. **(C)** KIF19 expression in glia declines as sample autolysis time increases. **(D,E)** Example of gender and cell-type specific genes in glia **(D)** and granule cells **(E)**. Expression of PRKY in **(E)** is highest in granule cells, at intermediate levels in glia, and low in basket cells. **(F)** Table Showing the number of genes that are in common between eight groups of differentially expressed aging genes. No overlaps were found between age up-regulated genes from any cell type and down-regulated genes from any other cell type.

## SUPPLEMENTAL TABLES

**Table S1.** Summary of all RNA-seq datasets, including information about animals and sorts, related to methods

**Table S2.** Clinical information for all human tissue donors, related to methods, Figures 6 and S6.

**Table S3.** Specificity index calculations for mouse, rat, and human cell types using either species-specific annotations or with mouse-rat-human orthologous gene annotations, related to Figures 2 and S4.

**Table S4.** Differential expression analysis results for mouse and human-enriched genes, related to Figures 3, S5, and S6.

**Table S5.** Differential expression analysis results for the influence of clinical factors on gene expression in human samples, related to Figures 6 and S7.

**Table S6.** Full results from all gene ontology (GO) analyses performed in the paper, related to Figures 6.

